# Arginine-enriched mixed-charge domains provide cohesion for nuclear speckle condensation

**DOI:** 10.1101/771592

**Authors:** Jamie A. Greig, Tu Anh Nguyen, Michelle Lee, Alex S. Holehouse, Ammon E. Posey, Rohit V. Pappu, Gregory Jedd

## Abstract

Low-complexity protein domains promote the formation of various biomolecular condensates. However, in many cases, the precise sequence features governing condensate formation and identity remain unclear. Here, we investigate the role of intrinsically disordered mixed-charge domains (MCDs) in nuclear speckle condensation. Proteins composed exclusively of arginine/aspartic-acid dipeptide repeats undergo length-dependent condensation and speckle incorporation. Substituting arginine with lysine in synthetic and natural speckle-associated MCDs abolishes these activities, identifying a key role for multivalent contacts through arginine’s guanidinium ion. MCDs can synergise with a speckle-associated RNA recognition motif to promote speckle specificity and residence. MCD behaviour is tuneable through net-charge: increasing negative charge abolishes condensation and speckle incorporation. By contrast, increasing positive charge through arginine leads to enhanced condensation, speckle enlargement, decreased splicing factor mobility, and defective mRNA export. Together, these results identify key sequence determinants of MCD-promoted speckle condensation, and link the speckle’s dynamic material properties with function in mRNA processing.

## Introduction

Eukaryotic cells simultaneously execute and coordinate a vast array of complex molecular reactions. These can be unfavorable, mutually incompatible, or require special environmental conditions that necessitate segregation and organization in subcellular organelles. Membrane-bound organelles attain their unique protein composition and environment by manipulating the permeability of their delimiting membrane. By contrast, ribonucleoprotein (RNP) bodies form through the condensation of constituent proteins and RNA (Banani et al., 2017; Shin and Brangwynne, 2017). Unlike membranous compartments, the constituents of these biomolecular condensates are in a dynamic equilibrium with the surrounding environment. As a result, membraneless organelles can readily form and dissolve in response to cellular and physiological cues (Brangwynne et al., 2009; Nott et al., 2015; Rai et al., 2018; Wippich et al., 2013). Well-established RNP bodies include nucleoli (Boisvert et al., 2007; Brangwynne et al., 2011; Feric et al., 2016), nuclear speckles (Spector and Lamond, 2011), paraspeckles (Fox et al., 2002), Cajal bodies (Cioce and Lamond, 2005; Sawyer et al., 2017) and promyelocytic leukemia bodies (Dyck et al., 1994; Lallemand-Breitenbach and de Thé, 2018) in the nucleus; and P granules (Brangwynne et al., 2009; Seydoux, 2018), stress granules (Kedersha et al., 1999; Protter and Parker, 2016), and P-bodies (Decker and Parker, 2012; Sheth and Parker, 2003) in the cytosol.

The formation of RNP bodies has been linked to liquid-liquid phase separation (LLPS) of constituents (Elbaum-Garfinkle et al., 2015; Feric et al., 2016; Molliex et al., 2015; Nott et al., 2015; Patel et al., 2015). Multivalent interactions among repetitive protein-protein or protein-RNA interaction domains / motifs are the key determinants of LLPS (Dennis, 2015; Fung et al., 2018). Multivalent contacts can be achieved with tandem repeats of folded domains (Li et al., 2012), through low-complexity (LC) intrinsically disordered regions (IDRs) (Halfmann, 2016; Lin et al., 2015; Mittag and Parker, 2018), or a combination of the two (Mitrea et al., 2016; Zhang et al., 2015). In these systems, individual contacts are relatively weak. However, with increasing valence, the combined effect of many weak interactions overcome the entropic cost of LLPS (Hyman et al., 2014). In many cases LC-IDRs have been shown to be necessary and sufficient for LLPS. This raises key questions regarding the types of residues that promote intermolecular contacts, and the extent to which these determine condensate identity. Emerging evidence identifies a key role for intermolecular contacts based on specifically patterned charged and aromatic residues (Brangwynne et al., 2015; Nott et al., 2015; Pak et al., 2016; Wang et al., 2018).

Nuclear speckles concentrate various factors involved in the regulation of gene expression, mRNA processing and export (Galganski et al., 2017; Spector and Lamond, 2011). Like other condensates, speckles are highly dynamic (Phair and Misteli, 2000), display droplet-like properties (Kim et al., 2019; Marzahn et al., 2016; Misteli et al., 1997), and have both core and peripheral components (Fei et al., 2017). In addition to protein components and nascent mRNAs, speckles also contain non-coding RNA that is thought to regulate splicing factor activity and alternative splicing (Carter et al., 1991; Hutchinson et al., 2007; Prasanth et al., 2010; Tripathi et al., 2010). Speckles appear to assemble in part through the concerted action of a variety of LC-IDRs. A number of these can be sufficient for nuclear speckle incorporation. Such domains include: phospho-regulated arginine-serine (RS) dipeptide repeats found in a number of splicing factors (Cáceres et al., 1997; Li and Bingham, 1991; Misteli et al., 1998), domains enriched in basic-acidic dipeptides (Bishof et al., 2018), and histidine rich tracts (Alvarez et al., 2003; Salichs et al., 2009). However, the overall molecular basis for how LC-IDRs interact to promote speckle formation remains unclear.

Here, we examine low-complexity mixed-charge domains (MCDs), highly enriched for positive and negatively charged amino acids. Synthetic sequences allow a systematic examination of the effects of composition and patterning on MCD activity. Our data show that multivalence and arginine-enrichment are key features governing the related activities of condensation and speckle incorporation. MCD activity is strengthened with increasing net-positive charge through arginine, and diminished with increasing negative charge. Speckle identity and residence appear to be determined by the combined effect of RNA recognition though folded domains and the cohesive forces provided by MCDs. Expression of arginine-rich MCDs with net-positive charge leads to enlargement of speckles, increased residence time of RS-splicing factors and defects in mRNA export. Taken together, these results identify key determinants governing the role of MCDs in nuclear speckle condensation. They further show how altering the speckle’s material properties can lead to dysfunction in mRNA processing.

## Results

### Mixed-charge domains (MCDs) undergo condensation and localize to nuclear speckles

We previously showed that an arginine and aspartic acid (RD)-enriched MCD from the fungal SPA-5 (SPA-5^MCD^) protein forms a viscoelastic hydrogel. In the fungus, this activity is associated with the formation of cytoplasmic plugs that gate cell-to-cell channels (Lai et al., 2012). To explore the role of such sequences in animal cells, we expressed SPA-5^MCD^ as an mGFP-fusion in HeLa cells (SPA-5^MCD^-mGFP). In this context, SPA-5^MCD^-mGFP accumulates in nuclear speckles as revealed by co-localization with the speckle marker serine/arginine-rich splicing factor 1 (SRSF1) (Figures 1A and S1A). SPA-5^MCD^ is enriched in RD dipeptide repeats. However, it also contains other residues that could contribute to its behavior (Figure S1B). To isolate the role of the RD repeats, we produced pure RD-MCDs of varying lengths (RD^20^, RD^30^, RD^40^, RD^50^, RD^60^) and expressed these as mGFP-fusions. RD^20^-mGFP is diffusely localized throughout the cytosol and nucleoplasm. By contrast, RD^30^-mGFP through RD^60^-mGFP increasingly localize to speckles (Figures 1A, S1C and S1D). Interestingly, unlike the shorter sequences, RD^60^-mGFP also forms rods outside speckles that exclude SRSF1 (Figures 1A and S1E). These data show that a polypeptide composed purely of RD dipeptide repeats can promote repeat number-dependent speckle incorporation.

**Figure 1.**
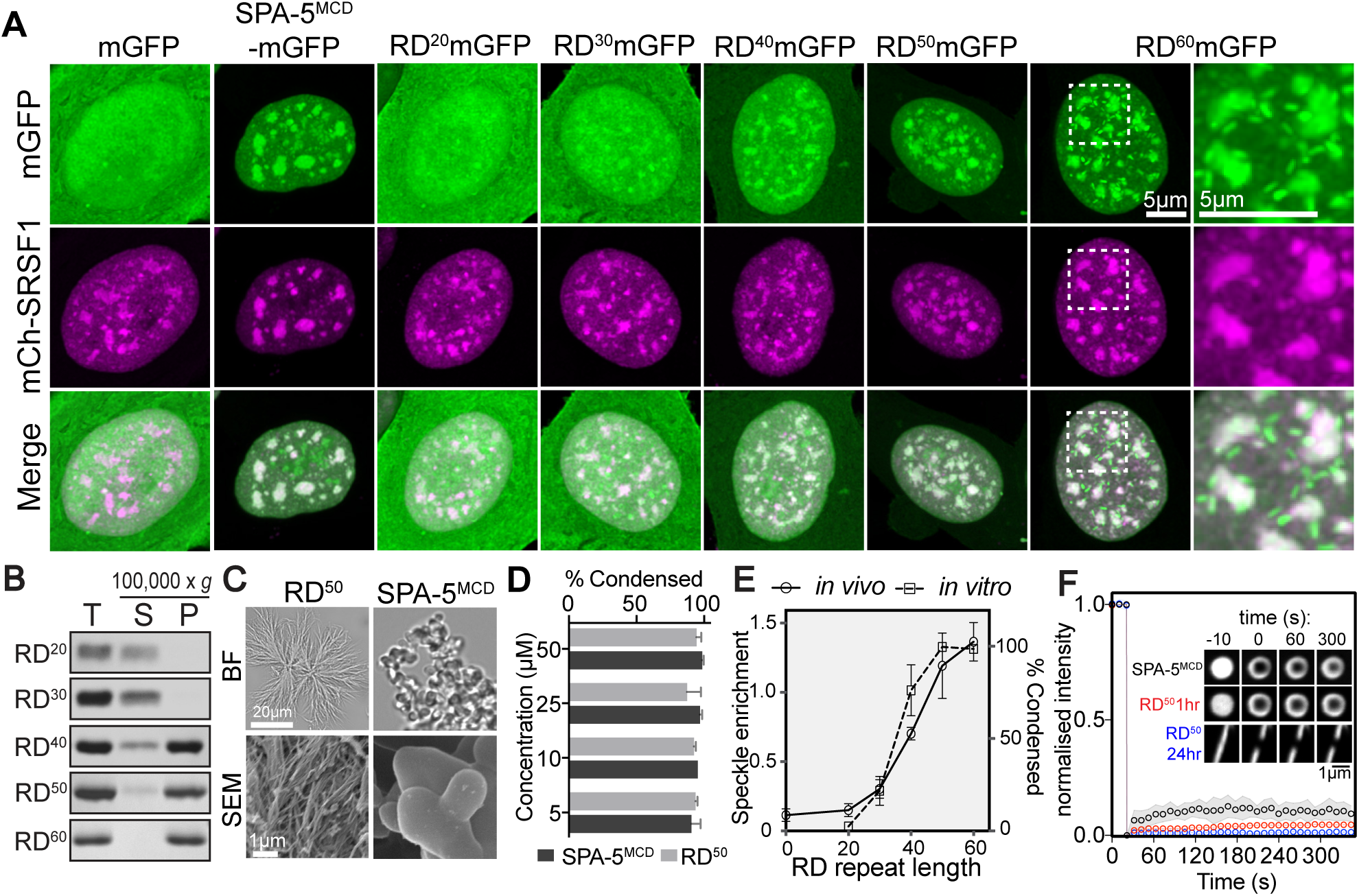
RD-dipeptide repeats associate with nuclear speckles and undergo condensation in a length-dependent manner. **A.** Representative images of nuclei in cells expressing the indicated mGFP-fusions. mCherry-SRSF1 identifies speckles. **B.** *in vitro* condensation of RD dipeptide length variants. Condensates are found in the pellet fraction following centrifugation at 100,000 x *g.* Total (T), supernatant (S) and pellet (P) fractions are shown. **C.** The appearance of SPA-5^MCD^ and RD^50^ condensates is shown by bright field (BF) (upper panels) and scanning electron microscopy (SEM) images (lower panels). **D.** Quantification of the percentage of total SPA-5^MCD^ and RD^50^ that pellet following condensation at the indicated protein concentrations. **E.** Quantification of speckle enrichment (*in vivo*) and condensation (*in vitro*) for the indicated RD variants reveals similar length-dependency. The degree of condensation is quantified as the fraction of pelleting material as compared to the total input. Average values are derived from three experiments as shown in part B. Speckle enrichment is the average speckle signal density divided by the average nucleoplasmic signal density, with the value of one set to zero. **F.** FRAP recovery curves for the indicated condensates. Insets show images of representative recoveries at the indicated time points. Figure S1 contains related information.

To examine the influence of RD repeat valence on the behavior of purified proteins, we produced the different length variants in *E. coli*. In this context, RD repeats form inclusion bodies, and were therefore purified under denaturing conditions before dialysis into a physiological buffer to allow condensation (see Materials and Methods, Figure S1F). Following centrifugation at 100,000 x *g*, RD^20^ and RD^30^ are found exclusively in the supernatant fraction (Figure 1B). By contrast, RD^40^ through RD^60^ progressively shift into the pellet fraction. The pelleting material consists of arrays of fibers that can be visualized using bright-field (BF) microscopy (Figures 1C and S1G). Scanning electron microscopy further shows that fibers are composed of bundles of fibrils (Figures 1C). By contrast, SPA-5^MCD^ assembles into multi-lobed structures. Both SPA-5^MCD^ and RD^50^ undergo condensation in the absence of molecular crowding agents, and assemble to completion at 5 μM (Figure 1D), indicating that condensation can occur at typical cellular protein concentrations (Hein et al., 2015). Quantification of *in vitro* assembly and *in vivo* speckle association reveals similar length dependency (Figure 1E). Thus, RD repeat condensation and speckle incorporation appear to be governed by the valence (number) of RD motifs.

To further examine MCD behavior, SPA-5^MCD^ and RD^50^ were produced as mGFP-fusions (Figure S1F). Proteins were purified in high salt, followed by dilution into a low salt buffer to allow condensation. In this assay, both mGFP-RD^50^ and mGFP-SPA-5^MCD^ condense into spherical droplets as observed by fluorescence and BF microscopy (Figure S1H). However, unlike the untagged proteins, this behavior requires the addition of a molecular crowding agent (Polyethylene glycol at 10 % (w/v)), suggesting that the mGFP moiety tends to counteract condensation. mGFP-RD^50^ droplets are metastable, maturing into rods over time. Fluorescence recovery after photobleaching (FRAP) of newly formed mGFP-RD^50^ and mGFP-SPA-5^MCD^ droplets is extremely limited, indicating that both condensates undergo an internal gelation transition (Figure 1F).

### Naturally occurring MCDs promote nuclear speckle assembly

To explore the functions of MCDs in mammalian cells, we searched the human proteome for sequences of greater than 60 amino acids in length with a fraction of charged residues (FCR) ⩾ 0.7 and a positive to negative charge ratio from 1:0.7 to 1:1.3. (Table S1). The output from this search is enriched for the speckle gene ontology (GO) term (Fold enrichment (FE) = 5.55, False discovery rate (FDR) = 8.89×10^-9^). Because a number of speckle proteins contain arginine-serine (RS) dipeptide repeats, which are regulated by phosphorylation (Colwill et al., 1996; Gui et al., 1994; Zhou and Fu, 2013), we also conducted the search including known phosphorylated serine residues as providers of negative charge (Table S1). This produces a further enrichment for the speckle GO term (FE = 7.63, FDR = 1.04×10^-27^). We next selected groups of arginine (R)- or lysine (K)-enriched MCDs (R-MCDs and K-MCDs) and examined their cellular localization as mGFP-fusions (Figures 2A, 2B, S2A-C and Table S2). R-MCDs from known nuclear speckle proteins, including the U1snRNP subunit U1-70k (snRNP70) (Bringmann and Lührmann, 1986; Verheijen et al., 1986) and the pre-mRNA cleavage and polyadenylation factor CPSF6 (Rüegsegger et al., 1998), are sufficient to promote speckle association. R-MCDs found in proteins that have not been previously localized to speckles are also sufficient for speckle incorporation. These include the polyadenylation factor FIP1L1 (Kaufmann et al., 2004), and the RNA polymerase II elongation regulator NELF-E (Yamaguchi et al., 1999). By contrast, K-MCDs from the nonsense mediated decay factor UPF2 (Lykke-Andersen et al., 2000) and the dentatorubral-pallidoluysian atrophy associated protein RERE (Yanagisawa et al., 2000) are diffusely distributed throughout the cell, while those from the chromatin remodelling factors NCOR1 and INO80 (Heinzel et al., 1997; Jin et al., 2005) appear to promote nucleolar localization (Figure 2B). The R-MCDs display varying tendencies towards condensate formation as pure proteins. mGFP-U1-70k and mCherry-CPSF6 R-MCDs form droplets, while the mCherry-NELF-E R-MCD forms fibrils. By contrast the mCherry-UPF2 K-MCD remains soluble under all conditions examined (Figures 2C and S2D). All the R-MCDs co-assemble with mGFP- or mCherry-RD^50^ (Figure 2D). Taken together, these data suggest that, under the conditions studied here, R-MCDs, but not K-MCDs form condensates. Their ability to co-assemble with RD^50^ further suggests that heterotypic R-MCD interactions contribute to nuclear speckle formation.

**Figure 2.**
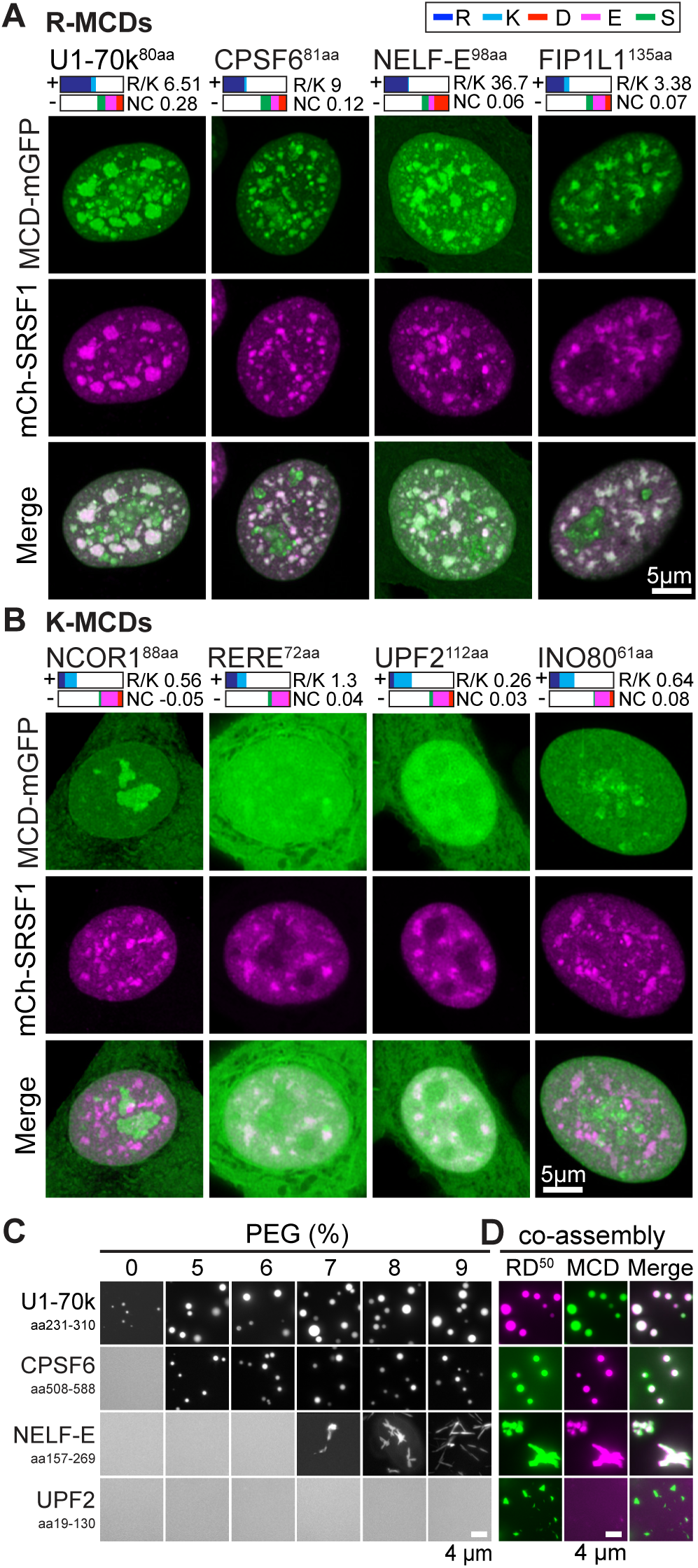
Naturally occurring R-MCDs associate with nuclear speckles and undergo phase separation as pure proteins. **A.** Images of representative nuclei for cells expressing the indicated R-MCD mGFP-fusions. mCherry-SRSF1 identifies speckles. Colocalization is shown in the Merge panel. For each protein, MCD length is identified in superscript. Colored boxes present the relative amino acid contribution of positively (+) and negatively (-) -charged amino acids and serine as identified in the legend. R/K ratio and net-charge (NC) are indicated. **B.** Images of representative nuclei for cells expressing the indicated K-MCD mGFP-fusions. mCherry-SRSF1 identifies speckles. Refer to A for legend description. **C.** *in vitro* phase separation of purified mGFP/mCherry-MCD fusions at the indicated PEG concentration. Condensates are identified by fluorescence microscopy. **D.** Co-assembly of MCDs from part C with mGFP/mCherry-RD^50^, identified as above. Figures S2A-D contain related information.

### Arginine plays a critical role in MCD condensation and nuclear speckle incorporation

We next sought to examine R-MCD function in the full-length spliceosome component U1-70k. U1-70k localizes to the speckles and foci corresponding to snRNP assembly bodies known as nuclear GEMs (Stejskalová and Staněk, 2014; Verheijen et al., 1986). In addition to the R-MCD examined above (MCD1), U1-70k contains a second smaller R-MCD (MCD2) (Figure 3A). Deletion of MCD1, but not MCD2 results in loss of speckle localization (Figures 3B and S2E). By contrast, R-MCD deletions do not affect GEM incorporation, which is known to rely on an N-terminal domain (Stejskalová and Staněk, 2014). To investigate the difference between the guanidinium moiety in R and the amine in K, we created a variant in which the MCD Rs are substituted with Ks. In the full-length protein, this change abolishes speckle incorporation but not assembly into GEMs. (Figures 3B and S2F). R to K substitution in MCD1-mGFP significantly diminishes its speckle incorporation while increasing localization to the nucleolus (Figures 3C, 3D and S2G). R to K substitution in MCD1-mGFP also abolishes its ability to undergo condensation in the *in vitro* assay (Figures 3E and S2D). Taken together, these data identify a key role for R in the U1-70k MCDs.

**Figure 3.**
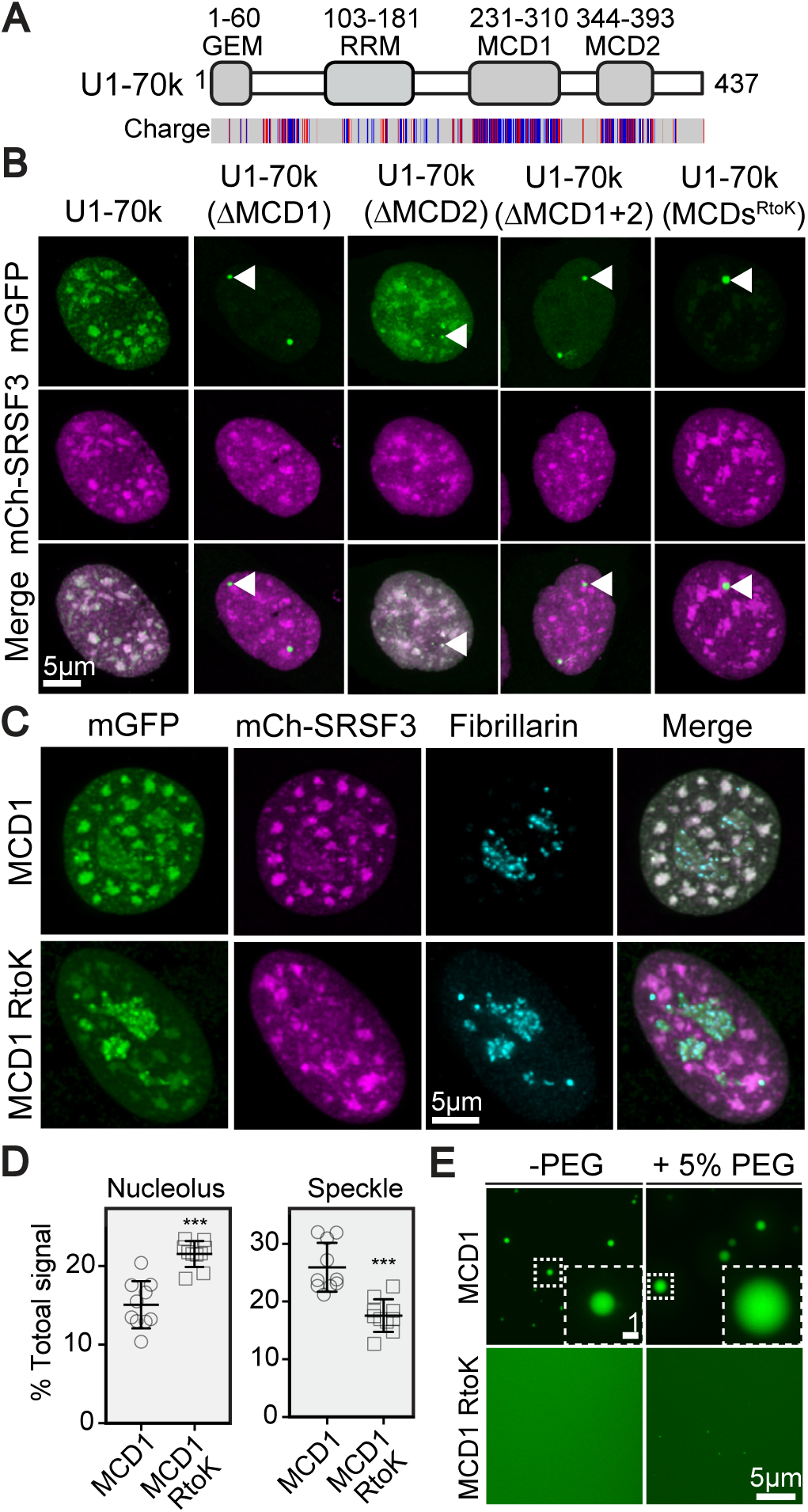
Arginine in U1-70k MCDs promotes nuclear speckle incorporation and condensate formation. **A.** Diagram of human U1-70k shows the position of the MCDs, RNA recognition motif (RRM), and GEM incorporation domain. The position of charged amino acids is shown in the lower panel. Blue: positively charged residues, Red: negatively charged residues. **B.** Images of representative nuclei from cells expressing the indicated U1-70k-mGFP variants. mCherry-SRSF3 identifies speckles. Arrowheads point to the GEM body. **C.** Images of representative nuclei from cells expressing the indicated mGFP-fusions. Fibrillarin staining identifies nucleoli. **D.** Quantification of the speckle and nucleolar signal as a fraction of the total nuclear signal for the mGFP-fusions shown in part C. *** p<0.0001. **E.** Fluorescence microscopy reveals the tendency of purified MCD1 and the MCD1 R to K variant-mGFP fusions to form condensates in the presence (+) and absence (-) of 5% PEG. The dashed box identifies the region that is shown magnified in the inset. Figures S2D-G contain related data.

Data presented thus far suggest that within MCDs, R and K have fundamentally different properties. To further investigate the role of different positive and negatively charged amino acids, we returned to pure synthetic dipeptide repeats and generated mGFP- and His-tagged versions of KD^50^, KE^50^, and RE^50^. Both K-MCDs, KD^50^ and KE^50^, fail to condense and do not localize to speckles (Figures 4A-B and S3A-B). This finding further corroborates the key role for the R guanidinium ion in MCD condensation and speckle incorporation. With respect to the choice of the negatively charged residue, RE^50^ undergoes condensation, but is not incorporated into speckles (Figures 4A-B). Instead, it forms punctate foci in the cytosol and nucleoplasm. As compared to pure RD repeat proteins, RE repeats condense at a lower critical length (Figures 4C and S3E-G) and associate weakly with speckles at intermediate lengths (Figures S3C-D). These data suggest that RE repeats favor self-association over the formation of heterotypic contacts that promote their incorporation into nuclear speckles. Unlike the fibrils formed by RD^50^, RE^50^ forms multi-lobed condensates *in vitro* (Figure 4D).

**Figure 4.**
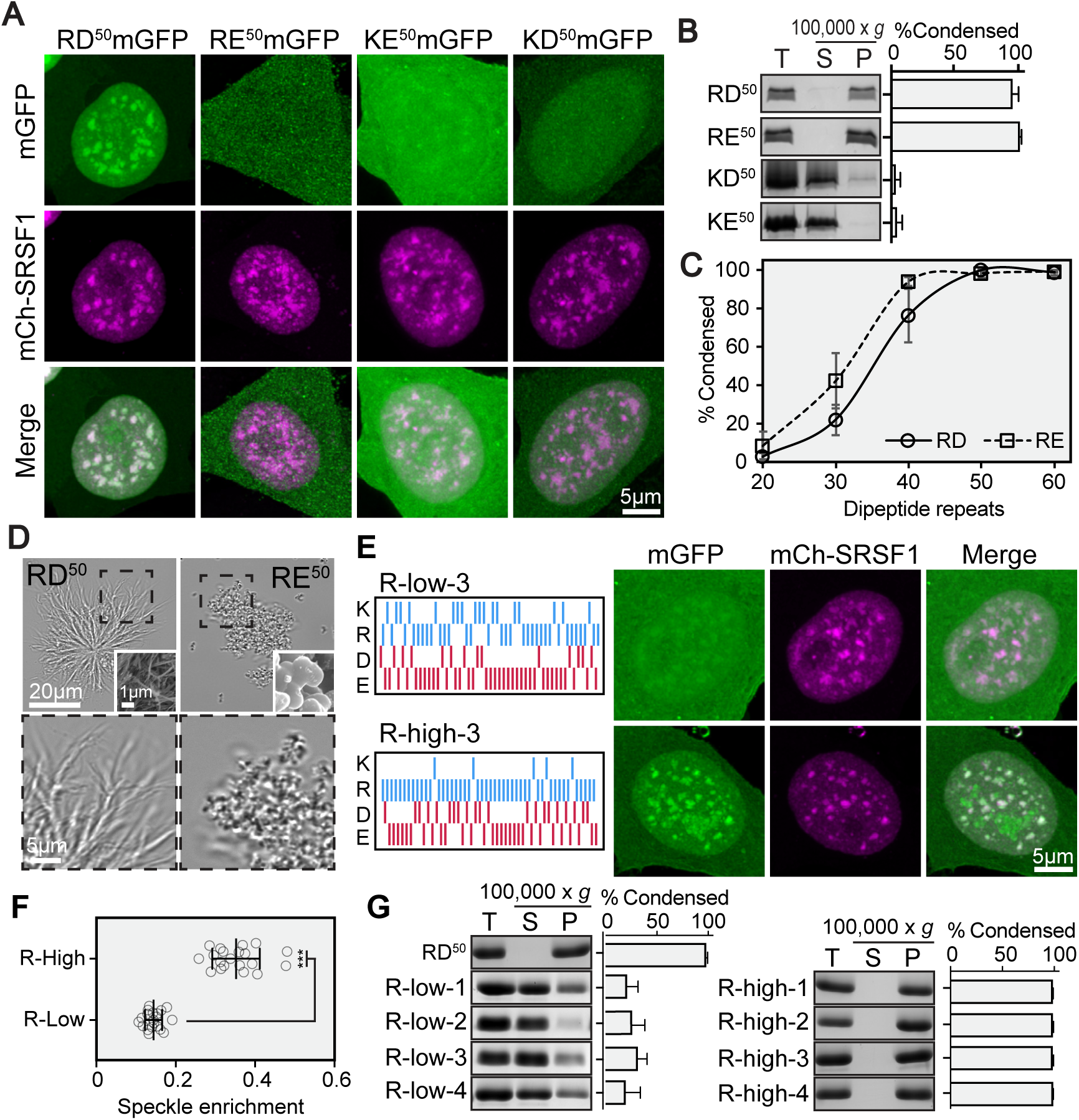
R is essential for condensation and nuclear speckle incorporation of MCDs. **A.** Images of representative nuclei from cells expressing the indicated mGFP-fusions. mCherry-SRSF1 identifies speckles. **B.** The indicated dipeptide repeat sequences were assayed for condensation as in Figure 1B. **C.** Quantification of length-dependent condensation for the indicated RD and RE repeat variants as in Figure 1E. **D.** BF images of the indicated condensates. Insets show SEM images. Dashed boxes indicate regions shown at higher magnification in the lower panels. **E.** Behaviour of randomized mixed-charge dipeptide repeat variants conforming to R-low and R-high R/K ratios. Diagrams on the left depict the position of the indicated amino acid residues. Images on the right are of representative nuclei from cells expressing the indicated R-low (upper panels) and R-high variant (lower panels). mCherry-SRSF1 identifies speckles. Figure S4 contains full analysis of four R-low and R-high variants. **F.** Quantification of speckle enrichment for the indicated sequence variants as in Figure 1E. *** p<0.0001. Data are derived from the full set of R-low and R-high variants. **G.** Quantification of *in vitro* condensation for the indicated MCD variants. Total (T), supernatant (S) and pellet (P) fractions are shown. Figures S3 and S4 contain all related data.

R-MCDs sufficient for speckle association (Figure 2A) (Bishof et al., 2018) have an average R/K ratio of 9 and an average E/D ratio of 1.38. To examine the behavior of dipeptide repeats composed of mixtures of R and K, and D and E we randomly generated four 100-residue dipeptide repeats that conform to either low (R/K=1.77) or high R content (R/K=9). All the sequences that have low R-content (R-low) exhibit weak speckle incorporation and condense poorly (Figures 4E-G and S4). By contrast, sequences that have high R-content (R-high) exhibit superior speckle recruitment and undergo condensation comparable to RD^50^ (Figures 4E-G and S4). These data corroborate findings regarding the R-enrichment of natural speckle MCDs, and suggest that MCDs generally require a high fraction of R for condensation and nuclear speckle incorporation. In the context of pure dipeptide repeats, RE^50^ behaves quite differently as compared to RD^50^ (Figures 4A-D). However, we note that when E is modestly enriched over D, as is observed in naturally occurring MCDs, randomly patterned mixed sequences show relatively uniform tendencies for both speckle incorporation and condensation (Figures 4E-G and S4).

### RS-domains appear to be phosphorylation-regulated MCDs

Arginine-serine (RS) domains are present in a variety of speckle-associated proteins (Jeong, 2017; Shepard and Hertel, 2009) (Table S3). RS domains can be sufficient to promote speckle incorporation (Cáceres et al., 1997), and are known to undergo extensive serine phosphorylation (Colwill et al., 1996; Gui et al., 1994; Zhou and Fu, 2013). Thus, RS domains appear to be R-MCDs whose net-charge can be tuned through phosphorylation. To investigate the behavior of isolated RS dipeptide repeats, we made synthetic RS sequences of 20-50 repeats. Upon expression as mGFP-fusions in HeLa cells, all these sequences undergo a degree of phosphorylation as revealed by phosphatase treatment (Figure S5A). RS^20^-mGFP is diffusely distributed through the nucleoplasm, and RS^30^ and RS^40^ form small foci in the nucleoplasm that do not appear to be associated with speckles (Figures S5B-C). By contrast, RS^50^-mGFP co-localizes with the speckle marker SRSF1. However, it also forms bright foci at the speckle periphery that exclude SRSF1 (Figure 5A). FRAP experiments demonstrate that RS^50^-mGFP bodies has extremely limited recovery on photobleaching compared with RD^50^-mGFP and the SRSF2 RS domain-mGFP, which are highly dynamic (Figures 5B and 5G). RD^50^ has a net-charge of zero. By contrast, unphosphorylated RS domains have a high net-positive charge, which drops progressively with increasing phosphorylation. Despite having fewer RS dipeptides (27), the RS domain from SRSF2 is more highly phosphorylated than RS^50^ (Figure 5C). Thus, RS domains appear to behave like phospho-tunable R-MCDs.

**Figure 5.**
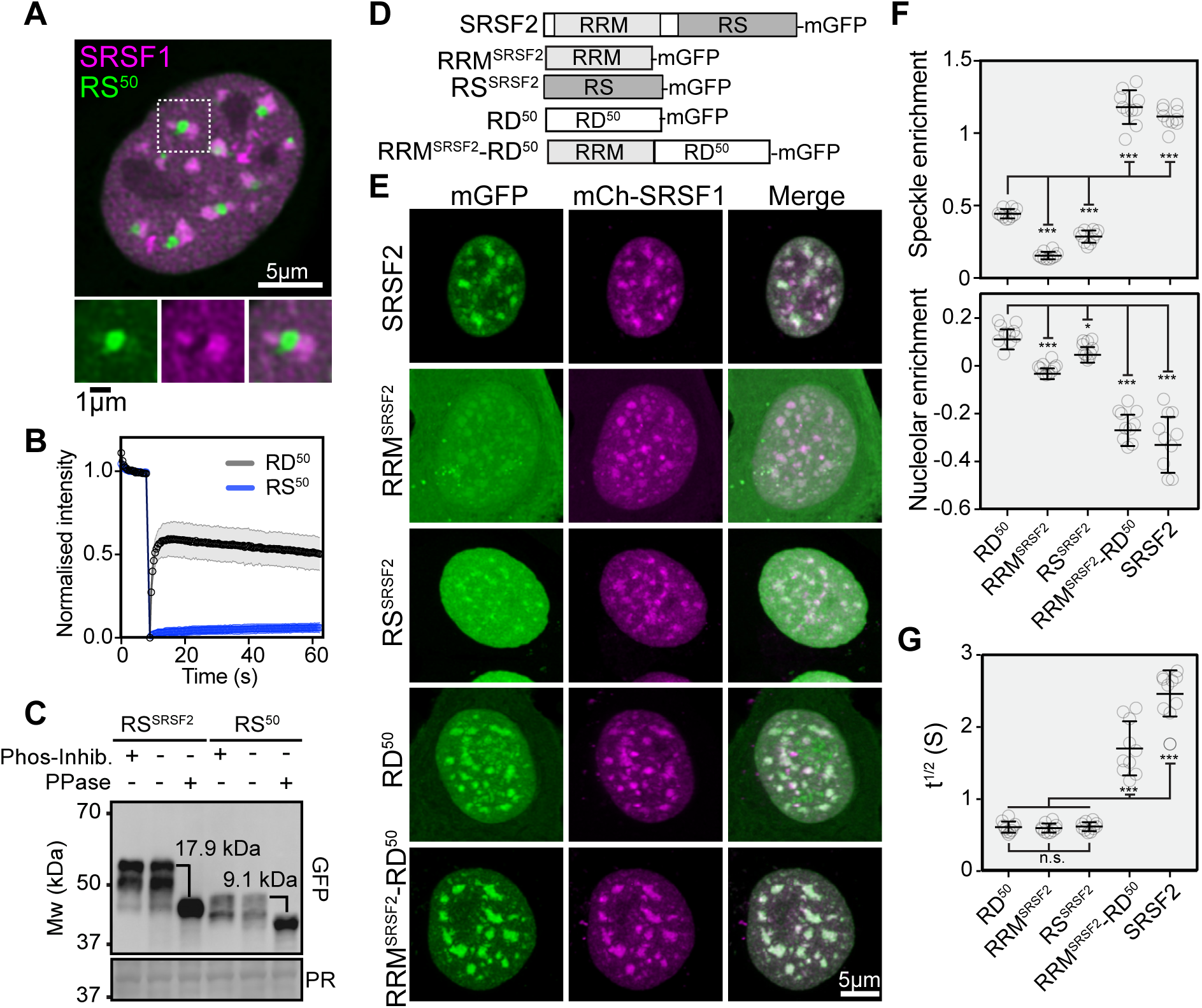
An RRM synergizes with an RS domain or R-MCD to specify nuclear speckle incorporation. **A.** Image of a representative nuclei in a cell expressing RS^50^-mGFP. Dashed box indicates the region magnified below. mCherry-SRSF1 identifies speckles. **B.** Limited FRAP recovery of RS^50^-mGFP nuclear bodies as compared to RD^50^-mGFP in speckles. **C.** Western blot comparing the migration of phosphatase (PPase) treated (+) and untreated (-) RS^50^-mGFP (RS^50^) with the RS domain from SRSF2 fused to mGFP (RS^SRSF2^). The difference in migration provides an estimate of the degree of phosphorylation, which is indicated in kDa. Ponceau red (PR) staining provides a loading control. **D.** Diagram showing the names and domain structure of the indicated mGFP-fusions. **E.** Images of representative nuclei from cells expressing the indicated mGFP-fusions. mCherry-SRSF1 identifies speckles. **F.** Quantification of the speckle-(upper panel) and nucleolar-enrichment (lower panel) of the indicated mGFP-fusions, as in Figure 1E. Nucleolar-enrichment is the average nucleolar signal density divided by the average nucleoplasmic signal density, with the value of one set to zero. **G.** FRAP of speckles labelled with the indicated mGFP-fusions. The half-times for recovery along with standard deviation are shown. For all quantification: *p<0.01, *** p<0.0001.

### Synergy between an R-MCD and RNA recognition motif enhance speckle specificity

A major question in the field of biomolecular condensates pertains to the molecular basis of condensate identity. The R-MCDs from U1-70K, CPSF6 and FIP1L1 exhibit varying degrees of incorporation into the nucleolus (Figure 2A). By contrast, full-length U1-70K (Figure 3B) (Verheijen et al., 1986) and CPSF6 (Cardinale et al., 2007) appear to be excluded from the nucleolus. RS-domain containing proteins and most of the naturally occurring R-MCD containing proteins identified here possess folded domains, among which RNA recognition motifs (RRMs) are the most common (Figure S2A) (Jeong, 2017; Shepard and Hertel, 2009). To better understand the interplay between R-MCDs and folded domains, we examined the behavior of the RNA recognition motif (RRM) from the splicing factor SRSF2 alone and in combination with RD^50^ (Figure 5D). As compared to full-length SRSF2, its RRM (RRM^SRSF2^) is weakly incorporated into speckles and has no tendency towards enrichment in the nucleolus (Cáceres et al., 1997) (Figures 5E-F). Interestingly, an RRM^SRSF2^-RD^50^ fusion protein is significantly more speckle-enriched than either domain alone, attaining a degree of enrichment comparable to full-length SRSF2 (Figures 5E-F). Moreover, the RRM^SRSF2^-RD^50^ fusion protein is excluded from the nucleolus to the same degree as full-length SRSF2 (Figures 5E-F). FRAP further reveals that the RRM^SRSF2^-RD^50^ fusion protein is more tightly associated with speckles than either RD^50^ or RRM^SRSF2^ alone (Figure 5G). Together, these findings indicate that RRMs and R-MCDs can work synergistically to promote speckle residency.

### Increasing MCD net positive charge through arginine enhances speckle cohesion leading to defects in mRNA export

RD^50^ has a net-charge of zero. However, naturally occurring R-MCDs tend to have a net-positive charge (Figure 2A), and this appears correlated to their incorporation into speckles and formation of condensates (Figures 2A and 2C). To examine the effect of net-charge, we generated R- and K-MCDs that use a 1 to 1 mixture of D and E, and expressed these as mGFP-fusions (Figures 6A-B and S6A). Starting with the net neutral MCD, regularly spaced substitutions for oppositely charged residues allow the systematic variation of net-charge per residue from zero to +0.1, +0.2 (or –0.1, –0.2) (Figure 6A). Increasing net-positive charge in the R-MCDs progressively enhances speckle residency (Figure 6B). This is reflected in enlarged speckles (Figure 6C), an increasing speckle to nucleoplasm signal ratio (Figure 6D), and slower FRAP recovery (Figure 6E). By contrast, increasing the net-positive charge in K-MCDs has little effect on speckle incorporation, but instead leads to increased accumulation within the nucleolus (Figures 6B, 6D and 6F). Increasing the net-negative charge of the R-MCD abolishes speckle incorporation, leading to diffuse localization throughout the cell (Figure 6B). This suggests that negatively charged residues counteract the condensation-promoting effect of arginine. *In vitro* behavior further supports the idea that the speckle-residency of the R-MCD variants is directly related to their propensity to undergo condensation. The R-MCD variant with a net-charge per residue of -0.1 is unable to undergo condensation, even in low salt buffer. By contrast, the net-neutral R-MCD shows modest activity, while the most robust condensation and highest degree of salt-resistance is observed with the highest net-positive charge variant (Figures 6G and S6B).

**Figure 6.**
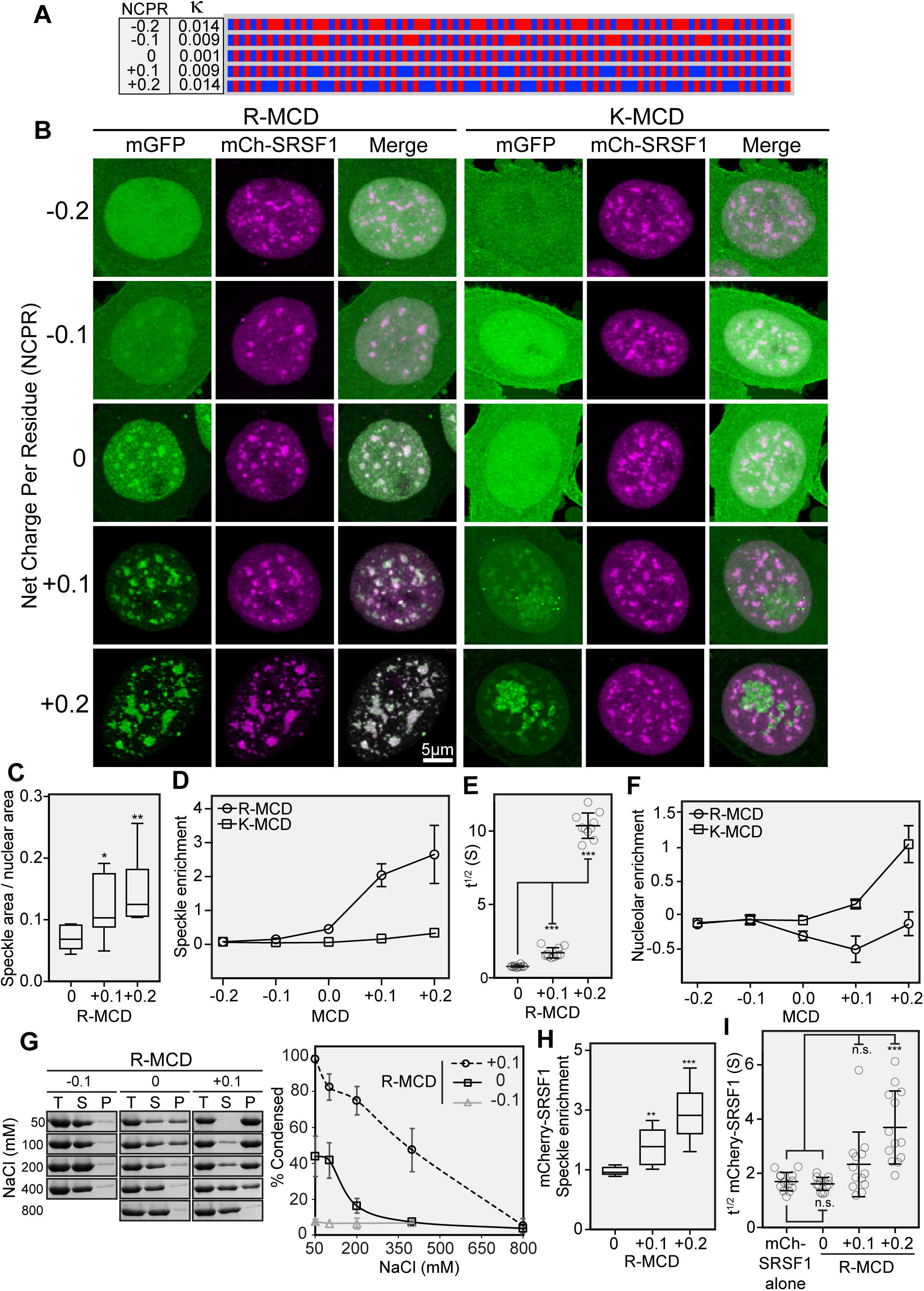
Increasing MCD net-positive charge through R enhances nuclear speckle cohesion. **A.** Diagram of the charge distribution in sequences designed to vary net-charge per residue (NCPR) between -0.2 and +0.2. Blue: positively charged residues, Red: negatively charged residues. The extent of charge mixing is indicated with the kappa (κ) value. **B.** Images of representative nuclei from cells expressing the indicated mGFP-fusions. mCherry-SRSF1 identifies speckles. **C.** Quantification of the fraction of nuclear area occupied by speckles for cells expressing the indicated R-MCD net-charge variants. **D.** Quantification of speckle enrichment for the indicated MCD net-charge variants, as in Figure 1E. **E.** Half-times of FRAP recovery for the indicated R-MCD net-charge variant labelled speckles. **F.** Quantification of nucleolar enrichment for the indicated net-charge variants, as in Figure 5F. **G.** *In vitro* condensation assays of the indicated R-MCD net-charge variants. Proteins were diluted to 10 µM at the indicated salt concentration and condensation was quantified by centrifugation as in Figure 1B. The left panels show SDS-PAGE analysis of a representative experiment. The graph to the right show’s quantification of the average values from three independent experiments. **H.** Speckle enrichment of mCherry-SRSF1 upon co-expression of the indicated R-MCD charge variants. Quantification as in Figure 1E. **I.** Half-times of FRAP recovery for mCherry-SRSF1 labelled speckles on co-expression of the indicated R-MCD. For all quantification: **p<0.001, *** p<0.0001. Figures S5 and S6 contain related data.

The patterning of charged residues has previously been shown to influence electrostatically-mediated phase separation and can be quantified by the parameter kappa (κ), which ranges between zero for fully mixed charges to one for fully segregated charges (Das and Pappu, 2013; Lin and Chan, 2017; Pak et al., 2016). Because net-charge per residue variants simultaneously vary net-charge and charge patterning, we produced net-neutral charge segregated RD variants in which repeats consisted of clusters of five (R5D5^10^) or seven (R7D7^7^) like-charged residues (Figure S6C). Segregating charge in this manner leads to a modest enhancement in speckle enrichment. However, a comparable nucleolar enrichment is also observed (Figures S6D-G). Thus, while the linear segregation of oppositely charged residues within the sequences of MCDs appears to promote condensation, the effect is not specific to speckles and its magnitude is small compared to that achieved by increasing net-charge through R.

We next examined the effect of R-MCD expression on the behavior of the RS-domain containing splicing factors SRSF1, SRFS2 and SRSF3. Expression of net-positive R-MCDs promotes speckle residency of all three (Figures 6B, 6H, S7A-B and S7D-E). Overall the magnitude of this effect increases with increasing net-positive charge. SRSF1 and SRSF3 display an increased speckle to nucleoplasm signal ratio upon expression of R-MCD^+0.1^. By contrast, a higher ratio is observed upon expression of R-MCD^+0.2^, and in this case, all three SRSFs are affected. Similarly, all three proteins show slower FRAP recovery with R-MCD^+0.2^ expression, while expression of R-MCD^+0.1^ only produces a significant effect on SRSF3 (Figures 6I, S7C and S7F). Together, these results show that R-MCDs can control the material properties of speckle condensates through heterotypic interactions that extend to RS-domain containing proteins.

To determine the functional impact of expressing high-net charge R-MCD variants, we used fluorescent in situ hybridization to localize total polyadenylated (Poly(A)) mRNA with Cy5-oligo-dT^30^ (Figure 7A). In cells expressing mGFP alone, or the net-neutral R-MCD, Cy5-oligo-dT^30^ signal is present in speckles and is uniform in the cytosol. In this group of transformants, there appears to be little variation in the overall staining pattern, which has a similar appearance in untransformed cells. By contrast, in cells expressing net-positive R-MCDs, Cy5-oligo-dT^30^ reveals significantly higher levels of poly(A) mRNA in speckles as compared to untransformed cells, or those expressing the net-neutral R-MCD (Figures 7A-B). Interestingly, accumulation of mRNA in the nucleus appears to be accompanied by a concomitant decrease in signal from the cytosol (Figure 7A). Moreover, the magnitude of this effect as measured by the ratio of average signal density in the nucleus to that in the cytosol correlates well with the expression level of net-positive R-MCD variants (Figure 7C). As with the effect on condensation and speckle size, the variant with the highest net-positive charge has the strongest effect. Together, these data show that increasing speckle cohesion results in the aberrant accumulation of mRNA within speckles. The loss of cytosolic mRNA signal further suggests significant defects in mRNA export.

**Figure 7.**
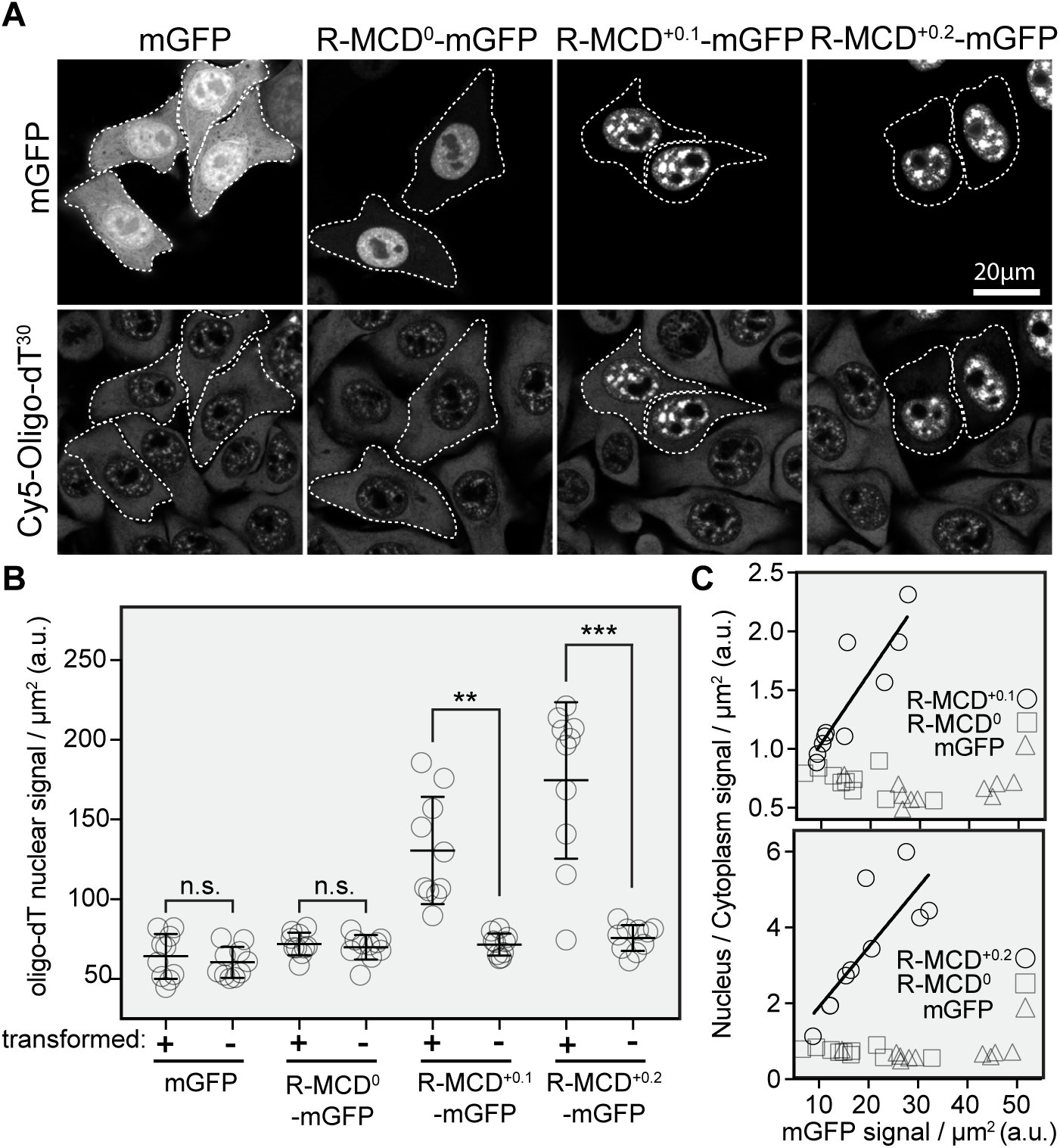
Increasing speckle cohesion leads to retention of polyadenylated mRNAs in the nucleus. **A.** Images of cells expressing mGFP alone or the indicated R-MCD charge variant fused to mGFP. A field of cells is shown to allow comparison between transformed and untransformed cells. Upper panels show mGFP fluorescence. Lower panes reveal staining of polyadenylated mRNA with Cy5-Oligo-dT^30^. The dotted white lines identify the periphery of transformed cells. **B.** Quantification of average density of nuclear polyadenylated mRNA upon expression of the indicated mGFP fusions. For each transformation, transformed (+) and untransformed (-) cells are compared. **C.** Quantification of the average density of nuclear to cytosolic mRNA upon expression of the indicated R-MCDs. The ratio is shown as a function of the level of R-MCD-mGFP fusion protein expression (x-axis). Values derived from expression of mGFP alone and the net-neutral R-MCD are shown on each graph for comparison.

## Discussion

Despite considerable progress in understanding the molecular basis of compartmentalization via biomolecular condensation (Gomes and Shorter, 2019; Langdon et al., 2018; Nott et al., 2015; Pak et al., 2016; Wang et al., 2018), important open questions remain. In particular, how LC-IDRs contribute to compartment assembly and identity remains unclear. Here, we examine the influence of length, composition and charge patterning on the behavior of intrinsically disordered mixed-charge domains (MCDs). Using synthetic and natural MCDs, we show that R-enrichment is a key feature underlying condensation and recruitment to nuclear speckles (Figures 1A-B, 2A, 2C, 3B-E, 4A-B, 4E-G and 6B-D). By contrast, K-enriched MCDs (K-MCDs) do not display a comparable propensity towards condensation (Figures 2C, 3E, 4B, and 4G), and tend to promote nucleolar uptake (Figures 2B, 3C, 6B and 6F). R and K are generally considered related through their positively charged side chains. However, data presented here reveal fundamental differences. The R guanidinium ion comprises three planar nitrogen groups; an arrangement that allows simultaneous formation of charge-charge, pi-pi and cation-pi contacts. By contrast, the K side chain amine is monovalent, leading to an inability to engage in pi-pi contacts, and a weak tendency to form cation-pi contacts as compared to the R sidechain (Armstrong et al., 2016; Chong et al., 2018; Gallivan and Dougherty, 1999; Vernon et al., 2018). Distinctive interactions of R with aromatic residues (Bogaert et al., 2018; Vernon et al., 2018; Wang et al., 2018) and RNA (Boeynaems et al., 2019) have also been demonstrated in other condensates. As shown here for MCDs, R to K substitution in the Nuage component Ddx4 (Vernon et al., 2018), and the paraspeckle/stress granule protein FUS (Wang et al., 2018), significantly diminishes their ability to form condensates. R to K substitution in *C9orf72* derived proline-R repeats results in the formation of less viscous condensates with RNA (Boeynaems et al., 2019). Thus, R appears to play a fundamental context-specific role to promote the formation of different biomolecular condensates.

Synthetic sequences provide a powerful tool to explore the relationship between sequence composition, patterning, and phase behavior of IDRs. For pure RD repeats, condensation and speckle incorporation activities respond similarly to increasing repeat number (Figures 1A-B and 1E). Interestingly, unlike shorter variants, RD^60^-mGFP forms rod-shaped assemblies in the nucleus that exclude a marker of nuclear speckles (Figures 1A and S1E). This suggests that at high valence, RD-repeat sequences begin to favor self-association over the heterotypic contacts that drive them into speckles. In other systems, R has been shown to contribute to condensate formation through interactions with aromatic residues (Bogaert et al., 2018; Nott et al., 2015; Wang et al., 2018). However, such interactions are not available in sequences composed exclusively of positive and negatively charged residues. Moreover, the inability of K to substitute for R, suggests that charge-charge interactions alone are insufficient to promote condensation (Figures 3E and 4B). Interestingly, recent work documents a water-stabilized like-charge complex between guanidinium ion pairs (Hebert and Russell, 2019). Thus, R-R contacts may also contribute to MCD condensation. Other candidates for stabilizing interactions involving R-residues are likely to involve the apparent “Y-aromaticity” of R-residues, and hydrogen bonding to backbone carbonyl groups (Borders et al., 1994; Chong et al., 2018). Resolving the relative contributions of these different bonds will require further experimental and theoretical work.

Expression of R-MCDs with a high net-positive charge through R leads to enlarged speckles and increased residence time of R-MCDs, as well as the speckle components SRSF1, SRSF2 and SRSF3 (Figures 6B-C, 6E, 6H-I and S7). These observations, combined with the ability of RD^50^ to co-assemble with naturally occurring R-MCDs *in vitro* (Figure 2D), are consistent with a major role for extended heterotypic R-MCD contacts in speckle condensation. Interestingly, recombinant SRSF1 and SRSF2 have recently been shown to form condensates, which can incorporate the RNA Polymerase II C-terminal domain (Guo et al., 2019). While the role of RS domains and phosphorylation in this behavior remains unclear, this finding provides additional evidence for the ability of SRSF proteins to engage in an extended interaction network. The dynamic nature of speckles is likely to be intimately associated with their function in mRNA processing and export (Misteli et al., 1997). Indeed, the expression of high net-positive charge R-MCDs also leads to the aberrant accumulation of polyadenylated mRNA in speckles and diminished levels in the cytosol (Figure 7). Thus, increased cohesion of the speckles proteinaceous constituents results in defective mRNA release or export. While the precise cause of this defects remains to be determined, these results clearly indicate that altering the material properties of the speckle condensate impairs normal function.

Increasing MCD negative charge abolishes speckle-incorporation and condensation (Figures 6B, 6D and 6G). In the context of RS repeats, phosphoserine appears to provide for a similar effect. Poorly phosphorylated pure RS repeats with high net-positive charge terminally self-associate into aberrant nuclear bodies (Figures 5A, 5C and S5A-B). By contrast, the more highly phosphorylated RS domain from SRSF2 is weakly speckle-incorporated and highly dynamic (Figures 5C and 5E-G). Previous work suggests that the degree of phosphorylation of RS-splicing factors is employed as a switch to regulate speckle association/assembly. Phosphorylation is required for speckle residence (Keshwani et al., 2015; Lai et al., 2001; Ngo et al., 2005). However, additional phosphorylation promotes exit from the speckle (Colwill et al., 1996; Misteli et al., 1998; Velazquez-Dones et al., 2005). Further to this, the DYRK3 kinase has recently been shown to disassembles speckles during mitosis (Rai et al., 2018). Together, these data support a model where R-MCD activity is determined by the balance between cohesion through enrichment for R and dissolution promoted by the negatively charged residue, which can be provided by D, E or is tuneable through phospho-serine.

R-MCDs identified here are found in proteins associated with diverse aspects of mRNA processing. These include the central splicing factor U1-70k, poly-adenylation factors CPSF6 and FIP1L1, and the RNA polymerase 2 regulator, NELF-E. Previous work showed that U1-70k assembles into Alzheimer disease aggregates (Diner et al., 2014), and coprecipitates with the splicing factors RBM25 and LUC7L3 through basic-acidic dipeptide-repeat domains, which share physicochemical properties with R-MCDs examined here (Bishof et al., 2018). These data, combined with data presented here, suggest that diverse proteins associated with mRNA processing employ R-MCDs to promote speckle-incorporation and residence. SPA-5 was originally identified by machine learning as one of a group of cytosolic LC-IDR containing proteins that undergo condensation to gate fungal cell-cell channels (Lai et al., 2012). The finding that related LC-IDRs participate in speckle condensation in animal cells suggests an interesting type of convergent evolution in which similar sequences were selected to form condensates with dramatically different functions.

Isolated R-MCDs are incorporated primarily into nuclear speckles. However, they also display varying degrees of weak nucleolar labelling (Figures 1A, 2A-B, 3C, 6B, 6F, S6D and S6F). This tendency appears to be related to high net-positive charge through K (Figures 3C, 6B and 6F), or increased charge segregation (Figures S6C-F). This finding is consistent with previous work showing that K-rich tracts are sufficient for nucleolar uptake (Scott et al., 2010). K-MCDs do not appear to have a strong tendency to undergo condensation (Figures 3E and 4B). Thus, nucleolar incorporation of these sequences may occur through complex coacervation with negatively-charged protein regions and/or RNA, as observed for other nucleolar constituents (Mitrea et al., 2018; White et al., 2019). LC sequences based on R-residues that lack negative charges, such as the GR/PR dipeptide repeats derived from *C9orf72* and RGG motifs found in RNA binding proteins, also associate with the nucleolus (Feric et al., 2016; Kwon et al., 2014; Lee et al., 2016). Thus, the mixed-charged context of the R-MCDs appears to be an important determinant of speckle-specific incorporation.

Many membraneless organelles are RNP granules that function in distinct aspects of RNA processing. Speckles (Shevtsov and Dundr, 2011), paraspeckles (Clemson et al., 2009; Naganuma et al., 2012) and nucleoli (Berry et al., 2015; Falahati et al., 2016) all appear to be seeded at sites where the RNAs they act upon are produced. Recent work showing that a variety of condensate-forming IDRs can synergize with an RRM domain in yeast P-body assembly (Protter et al., 2018) has led to the proposal that RNA recognition by RRMs acts to seed condensate formation, while LC-IDRs are generally promiscuous and provide binding energy to drive condensation. In nuclear speckles, R-MCDs preferentially incorporate into speckles over nucleoli (Figures 1A, 2A, 6B, 6D and 6F). Moreover, fusion of RD^50^ to the RRM from SRSF2 significantly enhances residence time and speckle-specific incorporation as compared to either domain alone (Figures 5D-G). Thus, the RRM and R-MCDs each appear to contribute to both cohesion and specificity. Together, these findings lead to a model in which stereospecific recognition of RNAs by RRMs and the unstructured interactions of R-MCDs synergize to tune speckle condensation, composition and dynamics.

## Acknowledgments

GJ, JAG, TAN and ML are supported by the Temasek Life Sciences Laboratory and Singapore Millennium Foundation. The contributions of ASH, AEP, and RVP were supported by funds from the US National Institutes of Health (5R01NS056114 and 1 R01NS089932), the Human Frontier Science Program (RGP0034/2017), and the St. Jude Research Collaborative on membraneless organelles. Partial support for the contributions of ASH come from the MOLSSI foundation.

## Author Contributions

JAG and GJ conceived the project. JAG, ML and AEP performed experiments. TAN and ASH performed bioinformatics analysis. All the authors analyzed the data. JAG and GJ wrote the manuscript with input from all the authors.

## Methods

### Gene synthesis and expression

Synthetic MCD sequences were codon randomized allowing equal probability for all possible codons, and then synthesized as *NdeI*-*BamHI* fragments in pUC57 (GenScript). For expression in HeLa cells, sequences were sub-cloned into a modified pEGFP-N1 vector (Clonetech) where the *NdeI* site in the CMV promoter was removed and the A206K mutation introduced in eGFP to produce monomeric (mGFP) (Zacharias et al., 2002). MCD-mGFP fusions were produced through standard molecular biological techniques. Accessions and amplified regions for these constructs can be found in Table S4. All domain deletions and mCherry-fusion proteins were produced by overlap extension PCR, with mCherry-fusions subsequently cloned into pcDNA3.1+ (ThermoFisher # V79020). For expression and purification from *E. coli*, sequences were sub-cloned into a modified pET15b vector (Novagen #69661-3CN) with a C-terminal Thrombin-6xHIS sequence. To purify proteins as N-terminal mCherry or mGFP-fusions, a pET15b vector containing mGFP- or mCherry-*NdeI*-*BamHI*-TEV-6xHIS was used.

### Mammalian cell culture

HeLa cells were cultured in Dulbecco’s modified eagle’s medium (DMEM) (ThermoFisher # 10566016) supplemented with penicillin streptomycin and glutamine (ThermoFisher #10378016). For transfection, cells were cultured in 8-well microscopy chamber slides (ThermoFisher #155411PK) or 24-well plates. Transient transfection was carried out using lipofectamine 3000 (ThermoFisher) following the manufacturer’s instructions. 500 ng of plasmid DNA was used for 24-well plate transfections and scaled appropriately for other culture dishes. Unless otherwise stated, cells were fixed 48 hours post transfection with 4% paraformaldehyde (Electron Microscopy Sciences #15713S) in PBS for 15 minutes at room temperature. For imaging, cells were mounted in 90% glycerol phosphate buffered saline (PBS).

### Immunoblotting

HeLa cells grown in 24-well plates were washed four times with one volume of ice-cold PBS, before incubating for on ice for 10 minutes in 60 µL RIPA buffer (50 mM Tris pH 7.4. 150 mM NaCl, 1 % Triton X-100, 0.1 % SDS and 0.1% sodium deoxycholate), supplemented with Halt^TM^ Protease and Phosphatase Inhibitor cocktail (ThermoFisher #78440). The cell lysate was centrifuged at 20,000 x *g* for 10 minutes and the supernatant was boiled in loading dye. Phosphatase treatment was carried out on boiled lysates using alkaline phosphatase (NEB #M0290) for at least one hour at 37°C. Unless otherwise stated, 10 µL of sample was run on a 10% SDS-PAGE gel. Proteins were transferred to nitrocellulose, which was subsequently blocked with 10% Blotting-grade blocker (BioRad #1706404) in Tris buffered saline (50 mM Tris pH 7.4, 150 mM NaCl) with 0.1% Tween-20 (TBS-T). Primary antibodies were incubated in 2% milk TBS-T for two hours at room temperature or overnight at 4°C. Mouse anti-GFP (Roche 11814460001) was diluted 1:1000. Blots were washed three times for five minutes with TBS-T at room temperature. Secondary antibody, sheep anti-mouse HRP conjugate (GE #NA931), was diluted 1:8000 in 2% milk TBS-T and incubated for one hour at room temperature. Blots were washed as above before incubating with ECL reagents and imaging on a BioRad ChemiDoc™ MP Imaging System. Imaging was done at maximum resolution with 2×2 pixel binning and exposure determined by the software.

### Computational identification of naturally occurring MCDs

Protein sequences from the reviewed human proteome (Uniprot accession AUP000005640) were searched for regions with a minimum FCR of 0.7 and positive/negative charge ratio of 1:1±0.3 using a window size of 60 amino acids. To consider phosphorylation, the same search was conducted, but with known phosphorylated serine residues (https://www.phosphosite.org) considered as contributors of negative charge. RS-domains were identified in an analogous manner, with a window size of 60 amino acids, a fraction of 0.6 of serine + arginine, and with a serine/arginine ratio of 1:1 ± 1. The script used for MCD identification can be found at https://github.com/Anh-01-10-0000/Jedd_lab. Cellular component enrichment analysis was performed using the tool provided by http://geneontology.org/ with accessions of proteins containing at least one MCD satisfying the criteria stated above. Functional annotations for RS-domain proteins were generated using PantherDB (Mi et al., 2019). MCD sequence features were analyzed using the package localCIDER (Holehouse et al., 2017). The script used to identify RS domains can be found at https://github.com/alexholehouse/rscode.

### Immunostaining

Transiently transfected cells were cultured for 48 hours before washing once with PBS then fixing with 2% paraformaldehyde, 4% sucrose in PBS for five minutes at room temperature. Fixation was quenched with 100 mM glycine in PBS. Cells were permeabilized with 0.25% Triton X-100 in PBS for 10 minutes before washing twice in PBS. Blocking was carried out with 5% BSA, 5% normal goat serum and 0.05% Triton X-100 in PBS at room temperature for at least one hour. Rabbit anti-Fibrillarin (Abcam, ab5821) was diluted to 1:1000 in blocking solution and was incubated overnight at 4°C in a humid chamber. Cells were washed three times for five minutes with PBS containing 0.05% Triton X-100. The secondary antibody, goat anti-rabbit Dylight-405 (Thermo #35551), was diluted 1:1000 in blocking buffer and incubated with the cells for one hour at room temperature in a humid chamber. Cells were washed as above and then mounted in 90% glycerol-PBS for imaging. Cells were imaged as stated above.

### Super-resolution Imaging

Stimulated emission depletion (STED) imaging of RD^60^-mGFP was carried out on a Leica SP8 inverted microscope fitted with a 100x objective of N.A. 1.4. The 592 nm STED laser was set at 60% power and the 488 nm white-light laser at 7% power with a gain of 45%. 10 z-sections were taken over 1 µm with 16-line averages and a pixel size of 20 nm. Subsequently images were deconvolved using Huygens Professional. The compressed z-stack of all 10 sections is shown in Figure S1E.

### Imaging fixed mammalian cells

Paraformaldehyde-fixed cells were imaged on a Leica SP8 inverted confocal microscope with a 63x oil-immersion objective of N.A. 1.4. Images were acquired at Nyquist sampling rates, with a 51 nm pixel size in 140 nm z-sections at a scan speed of 400 Hz with 4x line averaging. For mGFP imaging, the fluorophore was excited at 488 nm with 5% laser power and the detector set to 500-550 nm at maximum 50% gain. For mCherry Imaging, the fluorophore was excited at 587 nm and the detector set to 600-650 nm with 50% gain. For staining using dylight405 the fluorophore was excited at 405 nm with the detector set at 405-420 nm with 50% gain. For Cy5-oligo-dT^30^ staining the fluorophore was excited at 631 nm with the detector set at 650-700 nm at 40% gain. Images of nuclei are compressed z-stacks covering the entire nucleus deconvolved using Huygens Professional (Scientific Volume Imaging) and processed using ImageJ (https://imagej.nih.gov/ij/). For Figures S6A, S6D and 7A, single medial sections are shown with 8x line averaging. For all quantification, single medial sections at 8x line-averaging were taken from a minimum of 10 different cells.

### Fluorescence in-situ hybridization (FISH)

Transiently transfected cells were cultured for 48 hours before fixation with 4% paraformaldehyde in PBS for 15 minutes. Cells were permeabilized with 0.1% Triton X-100 in 2x saline sodium-citrate (SSC) buffer for 15 minutes, before washing with 1 M Tris pH 8.0. Blocking was carried out with 0.005% BSA, 1 mg/ml yeast tRNA (ThermoFisher #AM7119) in 2x SSC buffer. Cells were then washed once with 1 M Tris pH 8.0 prior to hybridization. For probe hybridization, 1 ng/ul Cy5-oligo-dT30 was incubated in the blocking buffer with the addition of 10% dextran sulphate and 25% formamide for one hour at 37°C in a humid chamber. Cells were then washed twice with 4xSSC buffer and once with 2xSSC buffer before mounting in 90% glycerol in 2xSSC. Single medial planes were imaged with 8x line averaging using Nyquist sampling as above. Imaging settings were standardized on the +0.2 R-MCD expressing cells, with the same settings used for all other cells. For quantification, the nucleus and the cytosol were manually assigned from the oligo-dT signal.

### Quantification of speckle/nucleolar incorporation

The speckle or nucleolar enrichment is calculated as the average signal density in the speckle or nucleolus divided by the average density in the nucleoplasm. A 1:1 ratio is taken to indicate no enrichment and is set to zero for the graphs. Images of co-expressed mCherry-SRSF1 or Fibrillarin staining were thresholded to define a speckle or nucleolus mask, respectively. The periphery of the nucleus was manually assigned. An mCherry-SRSF1 mask was used to calculate the speckle area in Figure 6. In some cases, Nucleoli were defined through a nucleoplasmic void defined by the speckle marker. For Figure 3C, speckle and nucleolar signal are shown as a fraction of the total nuclear signal. For Figure 1 and S1, mCherry-SRSF1 signal was used to define speckles. five areas inside speckles and five areas in the nucleoplasm (total area of 1.055μm^2^) were used to generate an average signal density for each compartment. Enrichment was determined as defined above. 10 nuclei were analyzed for each transfection. All statistical analysis was carried out with a student’s t-test.

### Fluorescence recovery after photobleaching (FRAP)

For *in vivo* FRAP, HeLa cells were transiently transfected in 8-well chamber slides for 48 hours. Cells were then washed in pre-warmed DMEM without phenol red (Gibco) and incubated in this medium. Slides were incubated in a live-cell imaging (37°C, 5% CO_2_) stage top incubator (Tokai Hit) mounted on an Olympus FV3000 laser scanning confocal microscope fitted with a 60x lens of N.A. 1.3. 20 frames were taken at maximum acquisition speed for one-way scanning before nuclear speckles were bleached with 50% laser power for one frame, then 130 subsequent frames were taken. FRAP analysis was carried out using cellSens software (Olympus). Normalization was carried out against frames 5-15 with background subtracted from outside of the imaged cell. Half-times for recovery (t^1/2^) were calculated using the initial plateau of the recovery curve. Quantification was carried out from 10 separate experiments

For the FRAP of condensates formed *in vitro* (see below), proteins were diluted to 10 µM with a final buffer composition of 10 mM Tris-HCl pH 7.4, 150 mM NaCl and 1 mg/ml BSA, before addition of PEG to a final concentration of 10%, unless otherwise stated. Assemblies were left to form for one hour at room-temperature before imaging on an Olympus FV3000 inverted microscope at 100x magnification N.A. of 1.4. Three pre-bleach frames were taken before samples were bleached with 10% laser power at maximum speed for one frame before following the recover every 10 seconds for up to five minutes. The FRAP data were normalized as for the *in vivo* experiments, with quantification taken from five separate experiments.

### Denaturing purification of MCDs

MCD sequences were expressed as C-terminal Thrombin-6xHIS fusions in BL21 DE3 *E. coli*. Overnight grown cultures were diluted 1:20 into fresh LB medium at 30°C and expression was induced with 0.5 mM Isopropylthiogalactoside (IPTG) once the culture had reached an OD^600^ of 0.7. After four to five hours, cultures were harvested by centrifugation at 3,000 *x g* for 15 minutes at 4°C, washed once with ice-cold PBS and the pellets then frozen at -80°C. Pellets were re-suspended 8 M Urea, 100 mM NaH_2_PO_4_ 10 mM Imidazole, 10 mM Tris pH8.0, before 10 rounds of sonication on ice for 30 seconds at 30 W with 30 second intervals between. Insoluble material was pelleted at 18,000 *x g* for 20 minutes at 4°C. The supernatant fraction was incubated with HisPur Ni-NTA resin (Thermo #88223) in 20 mM Imidazole at 4°C for one hour with gentle agitation. The Ni-NTA resin was then washed three times with 10 bead volumes of lysis buffer containing 20 mM Imidazole, before elution in lysis buffer containing 350 mM imidazole. Samples were concentrated in three or 10 kDa molecular weight cutoff columns (Amicon #800010) before being flash frozen.

### Purification of mGFP/mCherry-fusion proteins

Sequences were expressed with an N-terminal mGFP/mCherry and a C-terminal TEV-6xHIS tag in BL21 DE3 *E. coli*. Stationary phase cultures were diluted 1:20 into fresh LB media at 28°C, and expression was induced with 0.5 mM IPTG once the culture had reached an OD^600^ of 0.7. Expression was carried out for 6-18 hours at 28°C agitating the culture at 175 rpm. Cells were harvested, washed once with ice-cold PBS and the pellets frozen at -80°C. Cell pellets were resuspended in approximately five volumes of lysis buffer (50 mM NaH_2_PO_4_ pH 8, 1 M NaCl, 10 mM Imidazole, 2 mM PMSF and protease inhibitor cocktail (Roche #4693132001). Cells were lysed by sonication on ice for 12 x 30 seconds at 30 W with one minute on ice in between. Insoluble material was pelleted at 18,000 *x g* for 20 minutes at 4°C. The supernatant fraction was incubated with HisPur Ni-NTA resin (ThermoFisher #88223) in 20 mM Imidazole at 4°C for 1 hour with gentle agitation. The bound resin was then washed 3 times with 10 volumes of lysis buffer containing 20 mM Imidazole, before elution in lysis buffer containing 350 mM imidazole. Eluted fractions were concentrated in a 10 kDa molecular weight cutoff column (Amicon #800010), and the 6xHIS tag was removed by TEV cleavage (AcTEV, ThermoFisher #12575015) overnight at 4°C. His-cleaved proteins were further purified by size-exclusion chromatography on a Superdex 75 10/300 (GE # 17517401) column attached to an AKTA Purifier fast protein liquid chromatography device (GE #29148721) in a 50 mM tris pH 7.4, 1 M NaCl buffer. Purified proteins were concentrated in this buffer and aliquots flash-frozen before storage at -80°C.

### *In vitro* phase separation assays of MCDs

Proteins were defrosted on ice and then diluted to the desired concentration in the denaturing lysis buffer. 100 µL samples were placed in Slide-A-Lyzer mini dialysis cups with a two kDa molecular weight cut-off (ThermoFisher #69553). Samples were dialyzed against a 10 mM Tris pH 7.4 buffer with 150 mM NaCl unless otherwise stated. Dialysis was left to proceed at room temperature for 24-48 hours before imaging or downstream sample processing. For imaging, condensates were gently pipetted to release them from the dialysis membrane and then visualized at 100x magnification using a DIC filter on an Olympus BX51 microscope fitted with a CoolsnapHQ camera (Photometrics) controlled by Metamorph. To determine the fraction of the protein that had condensed, the contents of the cups was gently resuspended by pipetting and transferred to a low-bind Eppendorf tube. 50 µL of sample was centrifuged at 100,000 x *g* for 30 minutes at 4°C. The supernatant was removed and boiled in loading dye. The pellet was resuspended in 50 µL of dialysis buffer before boiling in loading dye. Total post-dialysis (T), 100,000 x *g* supernatant (S) and 100,000 x *g* pellet (P) samples were run on a 15% SDS-PAGE gel. The relative amounts in each fraction were determined by staining with Instant*Blue* (Sigma #ISB1L). For quantitation, the gels were imaged with a LiCor odyssey scanner. To obtain the percentage of condensed material, the signal of band for the pellet fraction is represented as a percent of the total. The averages from three independent experiments are shown.

### *In vitro* phase separation assays: mGFP/mCherry-fusion proteins

Protein aliquots were defrosted on ice before centrifuging at 20,000 *x g* for 20 minutes at 4°C to pellet any aggregated material. Samples were diluted in low-bind Eppendorf tubes to 10 µM with a final buffer composition of 10 mM Tris pH 7.4, 150 mM NaCl, 1 mg/ml Bovine serum albumin (BSA). To simulate macromolecular crowding, polyethylene glycol (PEG) 3,350Da was added from a 50% stock to give the desired concentration. Samples were mixed well by gentle pipetting and incubated for one hour at room-temperature before imaging at 100x magnification on an Olympus BX51 Microscope. For production of figures, a maximum projection of five different areas is shown. For the quantification shown in Figure 6G the condensates formed by dilution were submitted to centrifugation at 100,000 x *g*, with the total, supernatant and pellet fractions being analyzed as described above. For co-assembly experiments (Figure 2D), both proteins were diluted to 10 µM, with the same final buffer composition as above. PEG 3,350 Da was added to 10% to simulate macromolecular crowding. Samples were well mixed and left for one hour at room temperature before imaging as above.

## Supplemental Information titles and legends

Please see supplemental figures

Tables with title and legends

Table S1. MCDs in the human proteome.

Table S2. Sequence parameters of MCDs studied in Figure 2.

Table S3. Predicted RS proteins and GO enrichment analysis.

Table S4. Plasmids and protein accessions.

