## Supplementary material for "Arginine-enriched mixed-charge domains provide cohesion for nuclear speckle condensation": Suplemental Figures

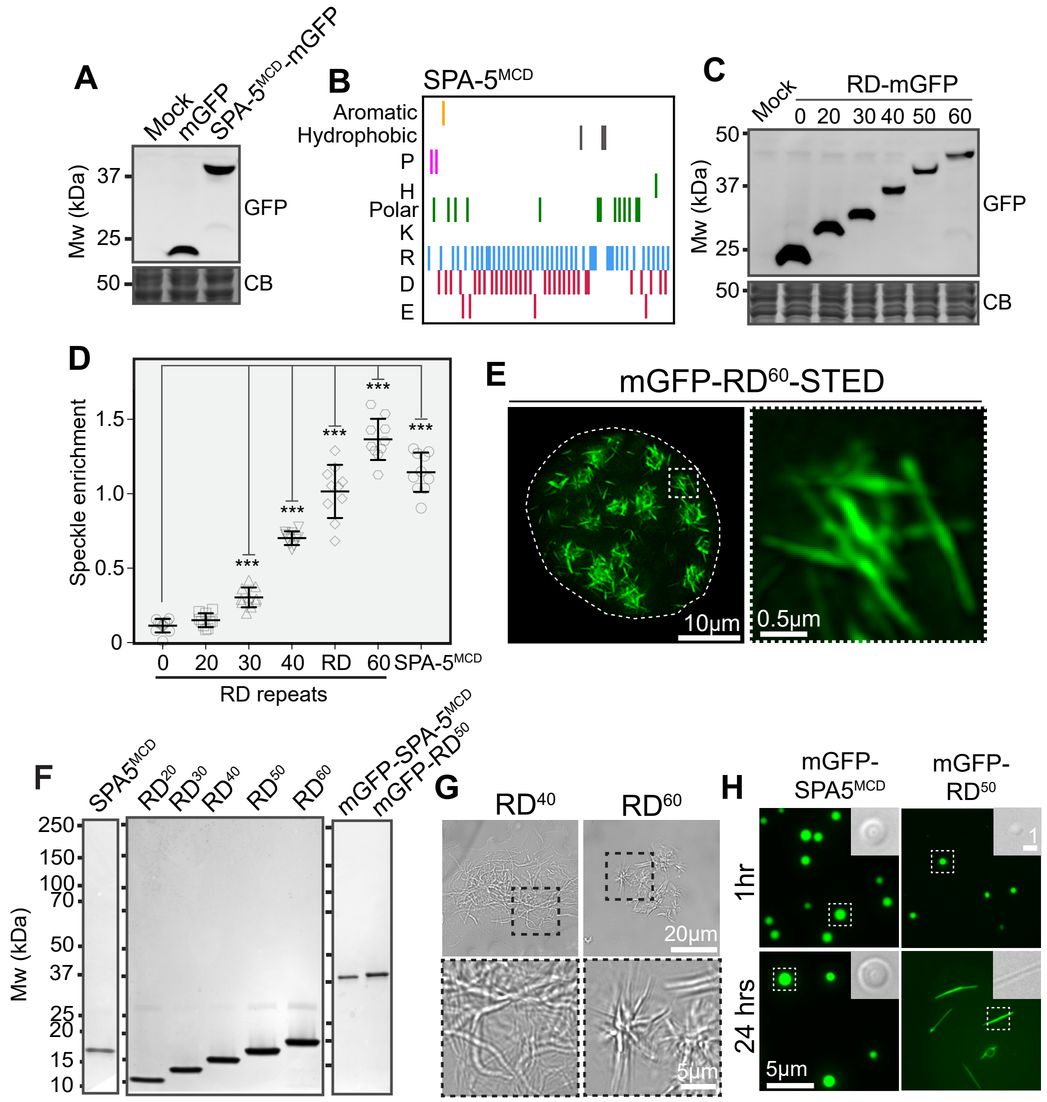


**Figure S1. Related to Figure 1. A.** Western blot showing SPA-5$^{MCD}$-mGFP expression in HeLa cells. **B.** Diagrammatic representation of SPA-5$^{MCD}$ composition. **C.** Western blot for the indicated RD dipeptide repeat length variant-mGFP fusions expressed in HeLa cells. Coomassie Blue (CB) staining provides a loading control. **D.** Quantification of the speckle enrichment (speckle/nucleoplasm average signal ratio) for the mGFP fusions shown in Figure 1A. These values are also shown in Figure 1E, compared with the *in vitro* condensation of the indicated sequences. ***$p<0.0001$. **E.** Stimulated emission depletion (STED) super-resolution imaging of mGFP-RD$^{60}$ nucleoplasmic fibrils. Area indicated by the dashed box is shown at higher magnification on the right. **F.** Coomassie stained gels of purified proteins used in Figures 1B-F. **G.** BF images of the condensates formed by the indicated RD-length variants. Dashed boxes define areas that are shown at higher magnification in lower panels. **H.** Fluorescence microscopy images of purified mGFP-SPA-5$^{MCD}$ and mGFP-RD$^{50}$ condensates at 1 and 24 hours (see Materials and Methods). Dashed boxes indicate regions shown in the inset by BF microscopy.

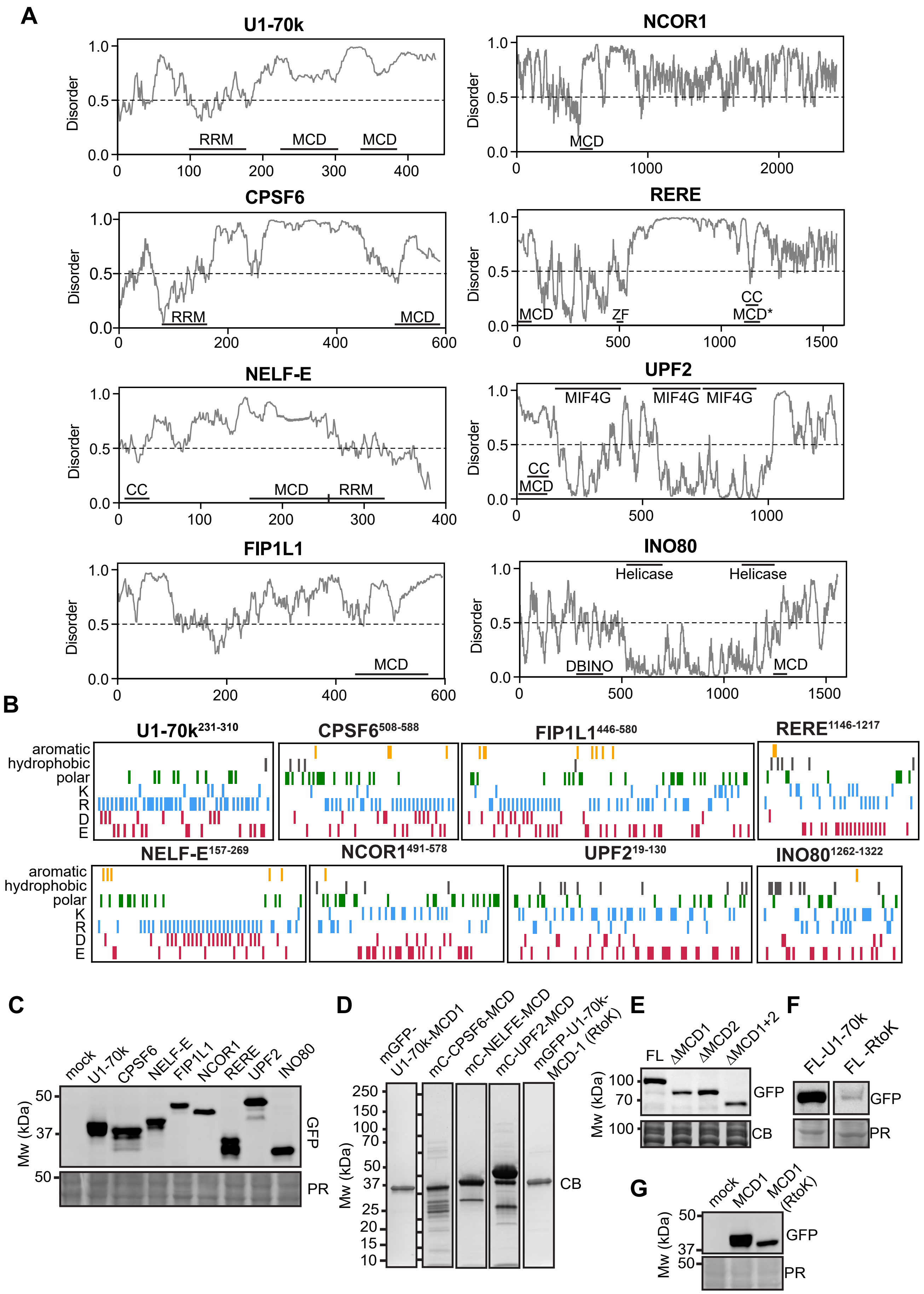
A
U1-70k
NCOR1
CPSF6
RERE
NELF-E
UPF2
FIP1L1
INO80
Disorder
RRM
MCD
MCD*
CC
ZF
MIF4G
Helicase
DBINO
B
U1-70k231-310
CPSF6508-588
FIP1L1446-580
RERE1146-1217
NELF-E157-269
NCOR1491-578
UPF219-130
INO801262-1322
aromatic
hydrophobic
polar
K
R
D
E
C
mock
U1-70k
CPSF6
NELF-E
FIP1L1
NCOR1
RERE
UPF2
INO80
Mw (kDa)
GFP
PR
D
mGFP-U1-70k-MCD1
mC-CPSF6-MCD
mC-NELFE-MCD
mC-UPF2-MCD
mGFP-U1-70k-MCD-1 (RtoK)
CB
E
FL
ΔMCD1
ΔMCD2
ΔMCD1+2
F
FL-U1-70k
FL -RtoK
G
mock
MCD1
MCD1 (RtoK)

**Figure S2. Related to Figures 2 and 3. A.** IUPRED Disorder predictions for proteins with MCDs used in Figure 2. MCDs as well as RNA recognition motifs (RRM), zinc fingers (ZF), coiled-coils (CC), helicase domains, the middle domain of eIF4G and the DBINO DNA binding domain are identified. **B.** Diagrams showing the residue composition and patterning of the MCDs used in Figure 2. **C.** Western blot of MCD-mGFP fusions expressed in HeLa cells in Figure 2. Ponceau red (PR) staining provides a loading control. **D.** Coomassie stained SDS-PAGE gels of purified proteins used in Figures 2C-D and 3E. **E.** Western blot of U1-70k deletion variants expressed in Figure 3B. FL = full-length U1-70k. Coomassie blue (CB) staining confirms equal loading **F.** Western blot of wild-type and R to K U1-70k variants shown in Figure 3B. **G.** Western blot of U1-70k MCD1-mGFP wild-type and R to K variant expressed in Figure 3C. PR staining confirms equal loading.

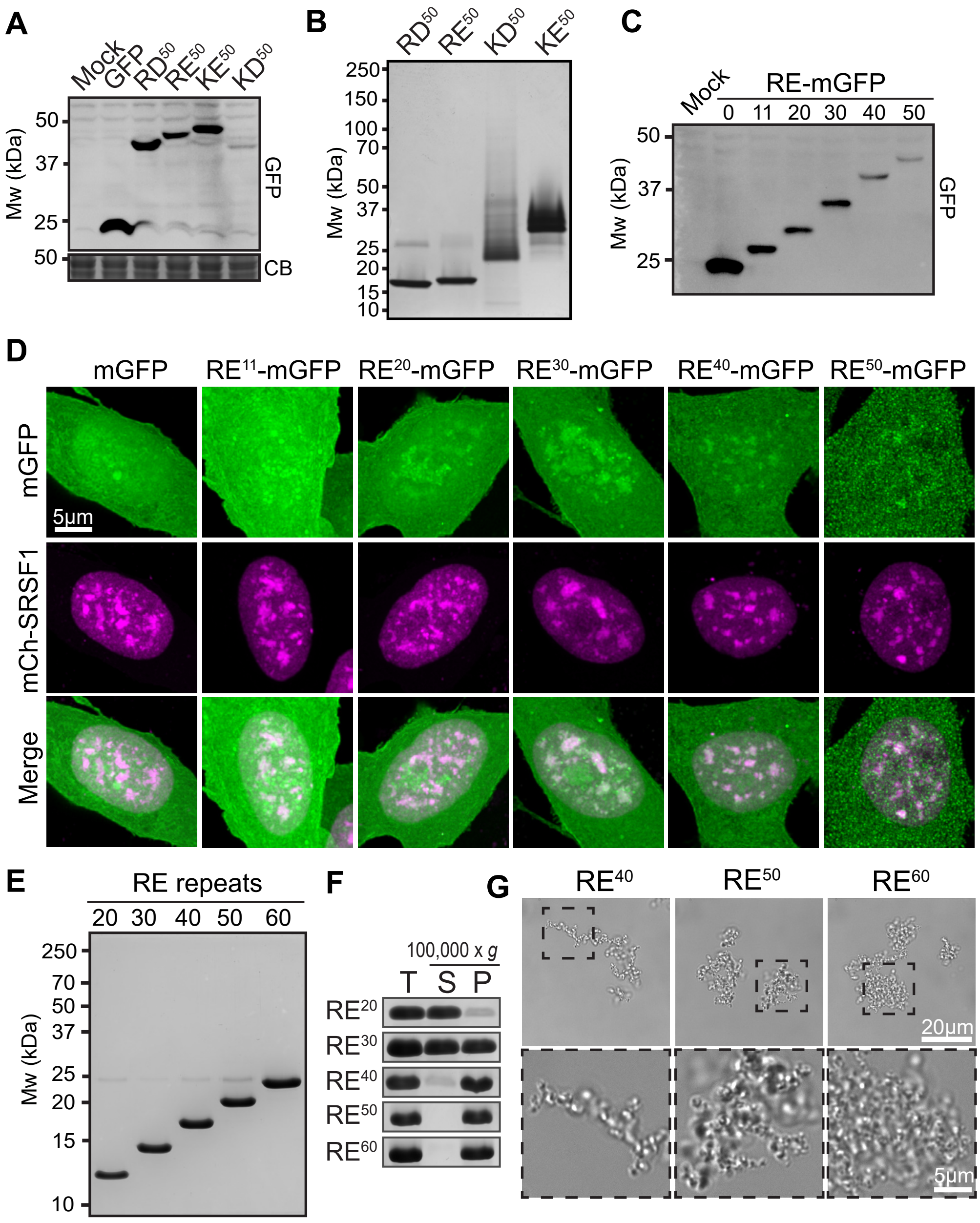


**Figure S3. Related to Figure 4. A.** Western blot of dipeptide-repeat mGFP-fusions expressed in HeLa cells in Figure 4A. **B.** Coomassie stained SDS-PAGE gel of purified dipeptide repeat variants used in Figures 4B-C. **C.** Western blot of RE length variant-mGFP fusions expressed in HeLa cells. **D.** Images of representative nuclei from cells expressing the indicated RE-repeat length variant-mGFP fusion. mCherry-SRSF1 identifies speckles. **E.** Coomassie stained SDS-PAGE gel showing purified RE dipeptide repeats used in Figures S3F-G. **F.** *In vitro* condensation of RE dipeptide repeat lengths, carried out as in Figure 1B - see Figure 4C for quantification. **G.** BF images of the condensates formed by the indicated RE dipeptide repeat-length variants. Dashed boxes designate regions that are shown at higher magnification in lower panels.

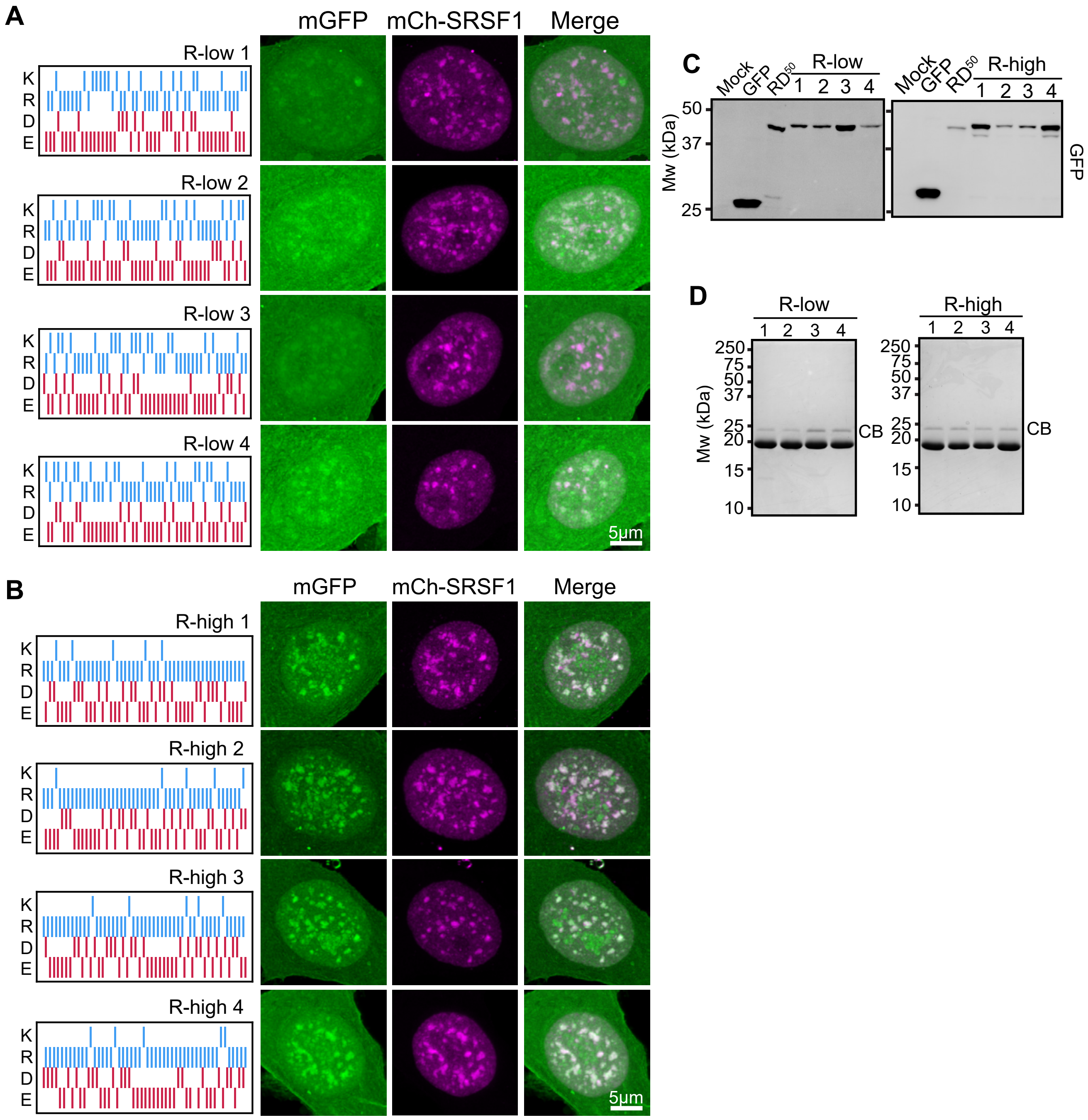


**Figure S4. Related to Figure 4. A.** Images of representative nuclei from HeLa cells expressing the indicated R-low MCD mGFP-fusions. Diagrams of the residue composition and patterning for the indicted MCDs are shown on the left. **B.** Images of representative nuclei from HeLa cells expressing the indicated R-high MCD mGFP-fusions. Diagrams of the residue composition and patterning for the indicted MCDs are shown on the left. **C.** Western blot shows expression of MCDs in parts A, and B. **D.** Coomassie stained SDS-PAGE gels of the indicated MCDs purified from *E. coli* and used in Figure 4G.

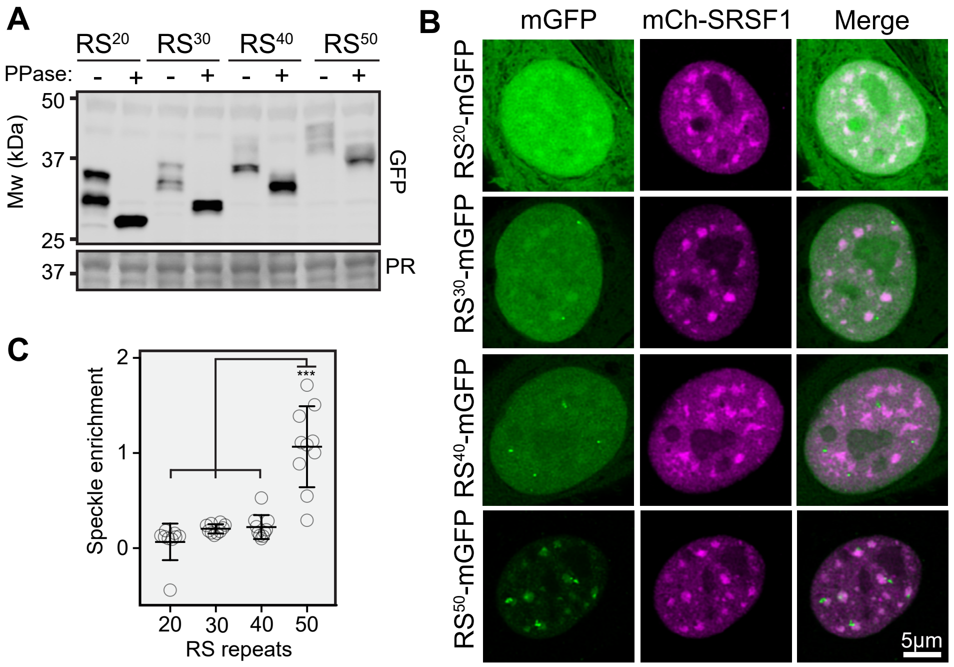


**Figure S5. Related to Figure 5. A.** Western blot of the indicated RS repeat mGFP-fusions expressed in HeLa cells. Phosphorylation is revealed by the difference in migration between phosphatase (PPase) treated (+) and untreated (-) samples. Ponceau Red (PR) staining provides a loading control. **B.** Representative images of nuclei from cells expressing the indicated RS dipeptide repeat mGFP-fusions. **C.** Speckle enrichment for the indicated RS repeat length variants. Quantification as in Figure 1E.

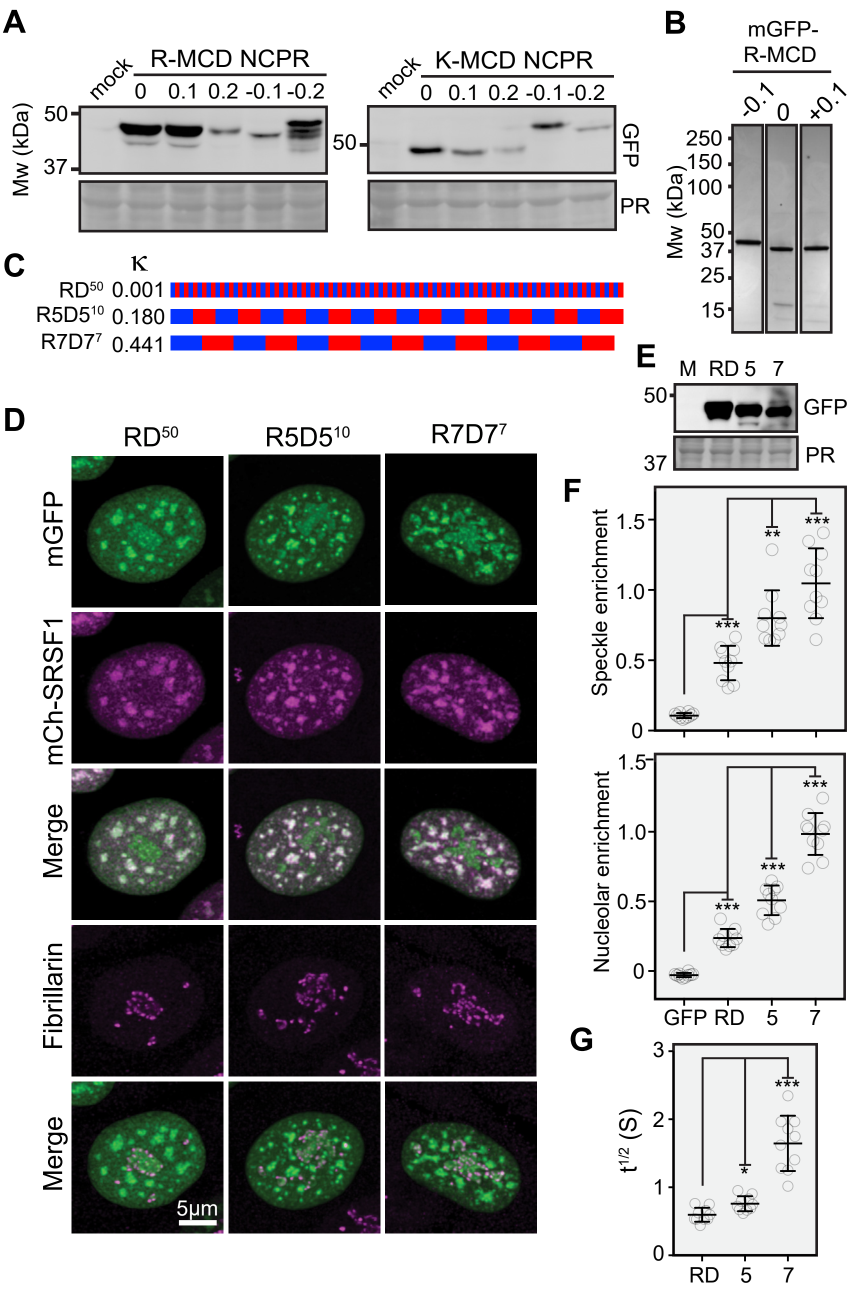


**Figure S6. Related to Figure 6. A.** Western blot showing the expression of the indicated R-MCD and K-MCD mGFP-fusions in Figure 6A. **B.** Coomassie stained gel of purified R-MCD net-charge variant mGFP-fusions. **C.** Diagram of the charge distribution for the indicated RD charge-mixing variants with their kappa (κ) value. Blue = R, Red = D. **D.** Representative nuclei from cells expressing the indicated RD charge-mixing variants as mGFP-fusions. mCherry-SRSF1 identifies speckles and Fibrillarin staining identifies the nucleolus. **E.** Western blot of indicated RD-mGFP variants expressed in part D. 5 = $R5D5^{10}$, 7 = $R7D7^{7}$. Ponceau red (PR) staining confirms equal loading. **F.** Speckle and nucleolar enrichment of the indicated RD-mGFP variants. Quantification as in Figures 1E and 5F, respectively **G.** Half-times for recovery of speckles labelled with the indicated RD-mGFP variants. *$p<0.01$, **$p<0.001$ *** $p<0.0001$.

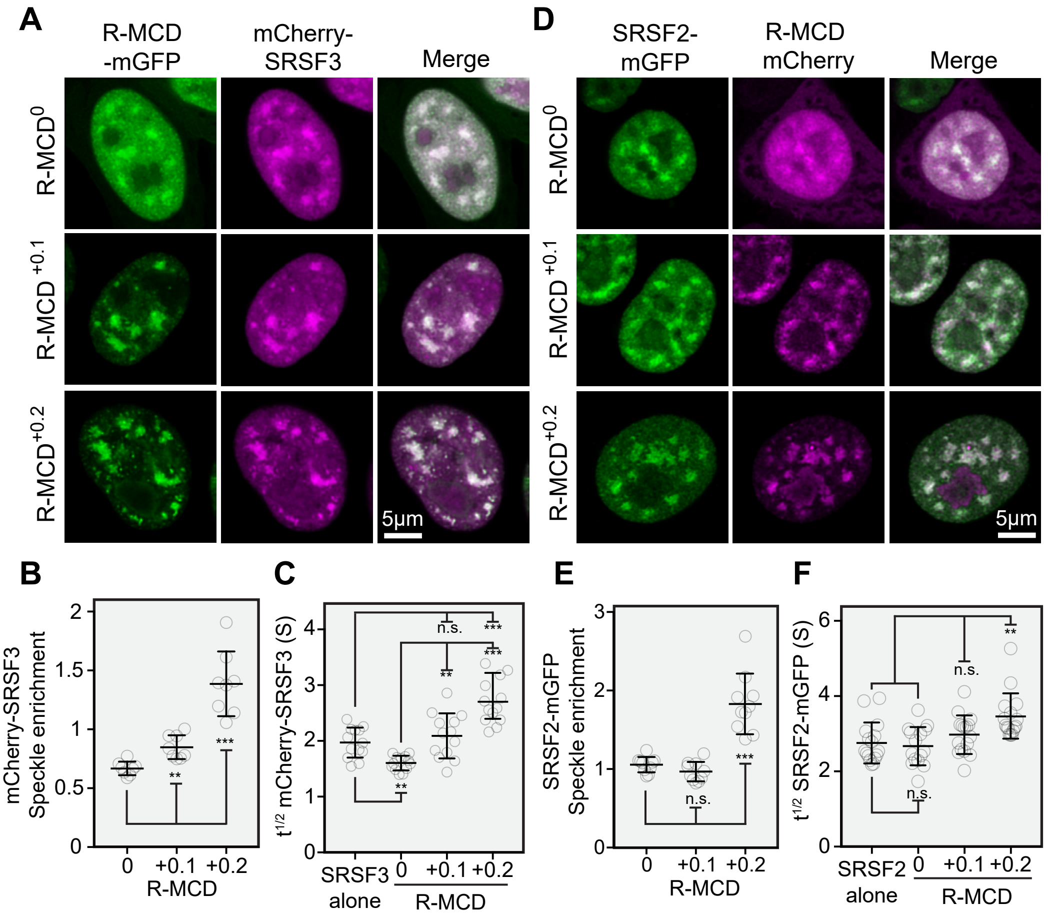


**Figure S7. Related to Figure 6. A.** Representative nuclei from HeLa cells co-expressing mCherry-SRSF3 with the indicated R-MCD net-charge variant. **B.** Speckle enrichment for mCherry-SRSF3 on co-expression of the indicated R-MCD net-charge variant. Quantification as in Figure 1E. **C.** Half-times for FRAP recovery of speckles labelled with mCherry-SRSF3 co-expressing the indicated R-MCD net-charge variant. **D.** Representative nuclei from HeLa cells co-expressing SRSF2-mGFP with the indicated R-MCD net-charge variant. **E.** Speckle enrichment for SRSF2-mGFP on co-expression of the indicated R-MCD net-charge variant. Quantification as in Figure 1E. **F.** Half-times for FRAP recovery of speckles labelled with SRSF2-mGFP co-expressing the indicated R-MCD net-charge variant. ** $p<0.001$ *** $p<0.0001$.
