## Supplementary material for "Arginine-enriched mixed-charge domains provide cohesion for nuclear speckle condensation": Table S2

### Table S4. Parameters of MCDs investigated in this study

| | N | f+ | f- | FCR | R:K | D:E | His | Tyr | NCPR | Kappa | Omega | Sigma | Delta | Hydropathy | Disorder Promoting |
|---|---|---|---|---|---|---|---|---|---|---|---|---|---|---|---|
| **U1-70k** | 80 | 0.563 | 0.288 | 0.850 | 0.867 | 0.391 | 0.000 | 0.000 | 0.275 | 0.084 | 0.585 | 0.089 | 0.059 | 0.978 | 0.988 |
| **CPSF6** | 81 | 0.370 | 0.250 | 0.617 | 0.900 | 0.500 | 0.074 | 0.049 | 0.123 | 0.059 | 0.252 | 0.025 | 0.034 | 1.684 | 0.901 |
| **NELF-E** | 98 | 0.378 | 0.316 | 0.694 | 0.973 | 0.710 | 0.020 | 0.010 | 0.061 | 0.037 | 0.557 | 0.005 | 0.025 | 1.538 | 0.939 |
| **FIP1L1** | 135 | 0.356 | 0.271 | 0.637 | 0.771 | 0.316 | 0.067 | 0.044 | 0.074 | 0.101 | 0.248 | 0.009 | 0.062 | 1.574 | 0.919 |
| **NCOR1** | 88 | 0.284 | 0.330 | 0.614 | 0.360 | 0.207 | 0.000 | 0.010 | -0.045 | 0.132 | 0.260 | 0.003 | 0.079 | 1.975 | 0.898 |
| **RERE** | 72 | 0.319 | 0.278 | 0.597 | 0.565 | 0.050 | 0.014 | 0.014 | 0.042 | 0.043 | 0.373 | 0.003 | 0.025 | 2.589 | 0.889 |
| **UPF2** | 112 | 0.348 | 0.321 | 0.670 | 0.205 | 0.222 | 0.009 | 0.000 | 0.027 | 0.061 | 0.298 | 0.001 | 0.040 | 2.027 | 0.893 |
| **INO80** | 61 | 0.377 | 0.295 | 0.672 | 0.391 | 0.222 | 0.000 | 1.000 | 0.082 | 0.169 | 0.229 | 0.010 | 0.110 | 2.292 | 0.803 |

**N**: Number of residues
**f-**: Fraction of negative residues
**f+**: Fraction of positive residues
**FCR**: Fraction of charged residues
**NCPR**: Net charge per residue
**Kappa**: κ (charge patterning parameter, discussed in the help section)
**Omega**: Ω (charge/proline patterning parameter, discussed in the help section)
**Sigma**: ⟨σ⟩, or the average sigma value for the entire sequence, where sigma quantifies the charge asymmetry.
**Delta**: The sequence's δ value represents the square deviation of every blob σ value from the sequence's mean σ value.
