## Supplementary material for "Arginine-enriched mixed-charge domains provide cohesion for nuclear speckle condensation": Table S4

### Table S4. Plasmids and Protein accessions.

#### Plasmids used in this study

| Name | Insert(s) |
|---|---|
| **Figure 1 and S1** | |
| pmEGFP-N1 | CMV(*ΔNdeI*)...*SalI*-*NdeI*-*BamHI*-GS-EGFP(A206K) |
| pcDNA3.1+ mCh-SRSF1 | CMV...*NheI*-mCherry-SRSF1-*XbaI* |
| pmEGFP-N1 SPA-5 | CMV(*ΔNdeI*)...*SalI*-*NdeI*-SPA-5(a.a.)-*BamHI*-GS-mEGFP |
| pmEGFP-N1 RD$^{20}$ | CMV(*ΔNdeI*)...*SalI*-*NdeI*-RD$^{20}$-*BamHI*-GS-mEGFP |
| pmEGFP-N1 RD$^{30}$ | CMV(*ΔNdeI*)...*SalI*-*NdeI*-RD$^{30}$-*BamHI*-GS-mEGFP |
| pmEGFP-N1 RD$^{40}$ | CMV(*ΔNdeI*)...*SalI*-*NdeI*-RD$^{40}$-*BamHI*-GS-mEGFP |
| pmEGFP-N1 RD$^{50}$ | CMV(*ΔNdeI*)...*SalI*-*NdeI*-RD$^{50}$-*BamHI*-GS-mEGFP |
| pmEGFP-N1 RD$^{60}$ | CMV(*ΔNdeI*)...*SalI*-*NdeI*-RD$^{60}$-*BamHI*-GS-mEGFP |
| pET15b mGFP | pT7-LacO-RBS-mGFP-*NdeI*-*BamHI*-TEV-6xHIS-HindIII |
| pET15b mGFP SPA-5 | mGFP-*NdeI*-SPA-5(a.a.)-*BamHI*-TEV-6xHIS-HindIII |
| pET15b mGFP RD$^{50}$ | mGFP-*NdeI*-RD$^{50}$-*BamHI*-TEV-6xHIS-HindIII |
| pET15b(cHIS) | pT7-LacO-RBS-*NdeI*-*BamHI*-Thrombin-6xHIS-*HindIII* |
| pET15b(cHIS) SPA-5$^{100}$ | *NdeI*-SPA-5(a.a.)-*BamHI*-Thrombin-6xHIS-*HindIII* |
| pET15b(cHIS) RD$^{20}$ | *NdeI*-RD$^{20}$-*BamHI*-Thrombin-6xHIS-*HindIII* |
| pET15b(cHIS) RD$^{30}$ | *NdeI*-RD$^{30}$-*BamHI*-Thrombin-6xHIS-*HindIII* |
| pET15b(cHIS) RD$^{40}$ | *NdeI*-RD$^{40}$-*BamHI*-Thrombin-6xHIS-*HindIII* |
| pET15b(cHIS) RD$^{50}$ | *NdeI*-RD$^{50}$-*BamHI*-Thrombin-6xHIS-*HindIII* |
| pET15b(cHIS) RD$^{60}$ | *NdeI*-RD$^{60}$-*BamHI*-Thrombin-6xHIS-*HindIII* |
| **Figure 2 and S2** | |
| pmEGFP-N1 U1-70k-PAT | CMV...*SalI*-*NdeI*-U1-70k(231-310)-*BamHI*-GS-mEGFP |
| pmEGFP-N1 CPSF6-PAT | CMV...*SalI*-*NdeI*-CPSF6(508-588)-*BamHI*-GS-mEGFP |
| pmEGFP-N1 NELF-E-PAT | CMV...*SalI*-*NdeI*-NELF-E(157-269)-*BamHI*-GS-mEGFP |
| pmEGFP-N1 FIP1L1-PAT | CMV...*SalI*-*NdeI*-FIP1L1(446-580)-*BamHI*-GS-mEGFP |
| pmEGFP-N1 NCOR1-PAT | CMV...*SalI*-*NdeI*-NCOR1(491-578)-*BamHI*-GS-mEGFP |
| pmEGFP-N1 RERE-PAT | CMV...*SalI*-*NdeI*-RERE(1146-1217)-*BamHI*-GS-mEGFP |

| | |
|---|---|
| pmEGFP-N1 UPF2-PAT | CMV...*SalI-NdeI*-UPF2(19-130)-*BamHI*-GS-mEGFP |
| pmEGFP-N1 INO80-PAT | CMV...*SalI-NdeI*-INO80(1262-1322)-*BamHI*-GS-mEGFP |
| pET15b mGFP-U170k-PAT | mGFP-*NdeI*-U1-70k(231-310)-*BamHI*-TEV-6xHIS |
| pET15b mCh-CPSF6-PAT | mCherry-*NdeI*-CPSF6(508-588)-*BamHI*-TEV-6xHIS |
| pET15b mCh-NELF-E-PAT | mCherry-*NdeI*-NELF-E(157-269)-*BamHI*-TEV-6xHIS |
| pET15b mCh-UPF2-PAT | mCherry-*NdeI*-UPF2(19-130)-*BamHI*-TEV-6xHIS |
| pET15b mCh-RD$^{50}$ | mCherry-*NdeI*-RD$^{50}$-*BamHI*-TEV-6xHIS |
| **Figure 3 and S2** | |
| pmEGFP-N1 *H.s.*U1-70k | CMV...*SalI*-U1-70k-*BamHI*-GS-mGFP |
| pmEGFP-N1 *H.s.*U1-70k(ΔP1) | CMV...*SalI*-U1-70k(Δ231-310)-*BamHI*-GS-mGFP |
| pmEGFP-N1 *H.s.*U1-70k(ΔP2) | CMV...*SalI*-U1-70k(Δ341-393)-*BamHI*-GS-mGFP |
| pmEGFP-N1 *H.s.*U1-70k(ΔP1+2) | CMV...*SalI*-U1-70k(Δ231-310+Δ341-393)-*BamHI*-GS-mGFP |
| pmEGFP-N1 *H.s.*U1-70k(P1+2 R to K) | CMV...*SalI*-U1-70k(231-310+341-393 RtoK)-*BamHI*-GS-mGFP |
| pcDNA3.1+ mCh-SRSF3 | CMV...*NheI*-mCherry-SRSF3-*XbaI* |
| pmEGFP-N1 U1-70k-P1 (R toK) | CMV...*SalI-NdeI*-U1-70k(a.a.231-310 RtoK)-*BamHI*-GS-mGFP |
| pET15b mGFP U1-70k-P1(R toK) | mGFP-*NdeI*-U1-70k(231-310 RtoK)-*BamHI*-TEV-6xHIS |
| **Figure 4, S3 and S4** | |
| pmEGFP-N1 RE$^{50}$ | CMV(*ΔNdeI*)...*SalI-NdeI*-RE$^{50}$-*BamHI*-GS-mEGFP |
| pmEGFP-N1 KE$^{50}$ | CMV(*ΔNdeI*)...*SalI-NdeI*-KE$^{50}$-*BamHI*-GS-mEGFP |
| pmEGFP-N1 KD$^{50}$ | CMV(*ΔNdeI*)...*SalI-NdeI*-KD$^{50}$-*BamHI*-GS-mEGFP |
| pET15b(cHIS) RE$^{50}$ | *NdeI*-RE$^{50}$-*BamHI*-Thrombin-6xHIS-*HindIII* |
| pET15b(cHIS) KE$^{50}$ | *NdeI*-KE$^{50}$-*BamHI*-Thrombin-6xHIS-*HindIII* |
| pET15b(cHIS) KD$^{50}$ | *NdeI*-KD$^{50}$-*BamHI*-Thrombin-6xHIS-*HindIII* |
| pmEGFP-N1 RE$^{20}$ | CMV(*ΔNdeI*)...*SalI-NdeI*-RE$^{20}$-*BamHI*-GS-mEGFP |
| pmEGFP-N1 RE$^{30}$ | CMV(*ΔNdeI*)...*SalI-NdeI*-RE$^{30}$-*BamHI*-GS-mEGFP |
| pmEGFP-N1 RE$^{40}$ | CMV(*ΔNdeI*)...*SalI-NdeI*-RE$^{40}$-*BamHI*-GS-mEGFP |

| | |
|---|---|
| pmEGFP-N1 $RE^{60}$ | CMV(*ΔNdeI*)...*SalI*-*NdeI*-$RE^{60}$-*BamHI*-GS-mEGFP |
| pET15b(cHIS) $RE^{20}$ | *NdeI*-$RE^{20}$-*BamHI*-Thrombin-6xHIS-*HindIII* |
| pET15b(cHIS) $RE^{30}$ | *NdeI*-$RE^{30}$-*BamHI*-Thrombin-6xHIS-*HindIII* |
| pET15b(cHIS) $RE^{40}$ | *NdeI*-$RE^{40}$-*BamHI*-Thrombin-6xHIS-*HindIII* |
| pET15b(cHIS) $RE^{60}$ | *NdeI*-$RE^{60}$-*BamHI*-Thrombin-6xHIS-*HindIII* |
| pmEGFP-N1 R-Low-1 | CMV(*ΔNdeI*)...*SalI*-*NdeI*-RKDE-$L1^{100aa}$ (R:K 1.56, D:E 1.63)-*BamHI*-GS-mEGFP |
| pmEGFP-N1 R-Low-2 | CMV(*ΔNdeI*)...*SalI*-*NdeI*-RKDE-$L2^{100aa}$ (R:K 1.56, D:E 1.63)-*BamHI*-GS-mEGFP |
| pmEGFP-N1 R-Low-3 | CMV(*ΔNdeI*)...*SalI*-*NdeI*-RKDE-$L3^{100aa}$ (R:K 1.56, D:E 1.63)-*BamHI*-GS-mEGFP |
| pmEGFP-N1 R-Low-4 | CMV(*ΔNdeI*)...*SalI*-*NdeI*-RKDE-$L4^{100aa}$ (R:K 1.56, D:E 1.63)-*BamHI*-GS-mEGFP |
| pmEGFP-N1 R-High-1 | CMV(*ΔNdeI*)...*SalI*-*NdeI*-RKDE-$H1^{100aa}$ (R:K 9, D:E 1.63)-*BamHI*-GS-mEGFP |
| pmEGFP-N1 R-High-2 | CMV(*ΔNdeI*)...*SalI*-*NdeI*-RKDE-$H2^{100aa}$ (R:K 9, D:E 1.63)-*BamHI*-GS-mEGFP |
| pmEGFP-N1 R-High-3 | CMV(*ΔNdeI*)...*SalI*-*NdeI*-RKDE-$H3^{100aa}$ (R:K 9, D:E 1.63)-*BamHI*-GS-mEGFP |
| pmEGFP-N1 R-High-4 | CMV(*ΔNdeI*)...*SalI*-*NdeI*-RKDE-$H4^{100aa}$ (R:K 9, D:E 1.63)-*BamHI*-GS-mEGFP |
| pET15b(cHIS) R-Low-1 | *NdeI*-RKDE-$L1^{100aa}$ (R:K 1.56, D:E 1.63)-*BamHI*-Thrombin-6xHIS-*HindIII* |
| pET15b(cHIS) R-Low-2 | *NdeI*-RKDE-$L2^{100aa}$ (R:K 1.56, D:E 1.63)-*BamHI*-Thrombin-6xHIS-*HindIII* |
| pET15b(cHIS) R-Low-3 | *NdeI*-RKDE-$L3^{100aa}$ (R:K 1.56, D:E 1.63)-*BamHI*-Thrombin-6xHIS-*HindIII* |
| pET15b(cHIS) R-Low-4 | *NdeI*-RKDE-$L4^{100aa}$ (R:K 1.56, D:E 1.63)-*BamHI*-Thrombin-6xHIS-*HindIII* |
| pET15b(cHIS) R-High-1 | *NdeI*-RKDE-$H1^{100aa}$ (R:K 9, D:E 1.63)-*BamHI*-Thrombin-6xHIS-*HindIII* |
| pET15b(cHIS) R-High-2 | *NdeI*-RKDE-$H2^{100aa}$ (R:K 9, D:E 1.63)-*BamHI*-Thrombin-6xHIS-*HindIII* |
| pET15b(cHIS) R-High-3 | *NdeI*-RKDE-$H3^{100aa}$ (R:K 9, D:E 1.63)-*BamHI*-Thrombin-6xHIS-*HindIII* |
| pET15b(cHIS) R-High-4 | *NdeI*-RKDE-$H4^{100aa}$ |

| | |
|---|---|
| | (R:K 9, D:E 1.63)-*BamHI*-Thrombin-6xHIS-*HindIII* |
| **Figure 5** | |
| pmEGFP-N1 SRSF2 | CMV(*ΔNdeI*)...*SalI*-*NdeI*-SRSF2-*BamHI*-GS-mEGFP |
| pmEGFP-N1 RRM$^{SRSF2}$ | CMV(*ΔNdeI*)...*SalI*-*NdeI*-SRSF2(aa9-101)-*BamHI*-GS-mEGFP |
| pmEGFP-N1 RS$^{SRSF2}$ | CMV(*ΔNdeI*)...*SalI*-*NdeI*-SRSF2(aa117-221)-*BamHI*-GS-mEGFP |
| pmEGFP-N1 RRM$^{SRSF2}$RD$^{50}$ | CMV(*ΔNdeI*)...*BglII*-SRSF2(aa9-101)-*NdeI*-RD$^{50}$-*BamHI*-GS-mEGFP |
| pmEGFP-N1 RS$^{20}$ | CMV(*ΔNdeI*)...*SalI*-*NdeI*-RS$^{20}$-*BamHI*-GS-mEGFP |
| pmEGFP-N1 RS$^{30}$ | CMV(*ΔNdeI*)...*SalI*-*NdeI*-RS$^{30}$-*BamHI*-GS-mEGFP |
| pmEGFP-N1 RS$^{40}$ | CMV(*ΔNdeI*)...*SalI*-*NdeI*-RS$^{40}$-*BamHI*-GS-mEGFP |
| pmEGFP-N1 RS$^{50}$ | CMV(*ΔNdeI*)...*SalI*-*NdeI*-RS$^{50}$-*BamHI*-GS-mEGFP |
| **Figure 6, S5, S6 and 7** | |
| pmEGFP-N1 R-MCD$^{0}$ | CMV(*ΔNdeI*)...*SalI*-*NdeI*-R-MCD$^{0}$-*BamHI*-GS-mEGFP |
| pmEGFP-N1 R-MCD$^{+0.1}$ | CMV(*ΔNdeI*)...*SalI*-*NdeI*-R-MCD$^{+0.1}$-*BamHI*-GS-mEGFP |
| pmEGFP-N1 R-MCD$^{+0.2}$ | CMV(*ΔNdeI*)...*SalI*-*NdeI*-R-MCD$^{+0.2}$-*BamHI*-GS-mEGFP |
| pmEGFP-N1 R-MCD$^{-0.1}$ | CMV(*ΔNdeI*)...*SalI*-*NdeI*-R-MCD$^{-0.1}$-*BamHI*-GS-mEGFP |
| pmEGFP-N1 R-MCD$^{-0.2}$ | CMV(*ΔNdeI*)...*SalI*-*NdeI*-R-MCD$^{-0.2}$-*BamHI*-GS-mEGFP |
| pmEGFP-N1 K-MCD$^{0}$ | CMV(*ΔNdeI*)...*SalI*-*NdeI*-K-MCD$^{0}$-*BamHI*-GS-mEGFP |
| pmEGFP-N1 K-MCD$^{+0.1}$ | CMV(*ΔNdeI*)...*SalI*-*NdeI*-K-MCD$^{+0.1}$-*BamHI*-GS-mEGFP |
| pmEGFP-N1 K-MCD$^{+0.2}$ | CMV(*ΔNdeI*)...*SalI*-*NdeI*-K-MCD$^{+0.2}$-*BamHI*-GS-mEGFP |
| pmEGFP-N1 K-MCD$^{-0.1}$ | CMV(*ΔNdeI*)...*SalI*-*NdeI*-K-MCD$^{-0.1}$-*BamHI*-GS-mEGFP |
| pmEGFP-N1 K-MCD$^{-0.2}$ | CMV(*ΔNdeI*)...*SalI*-*NdeI*-K-MCD$^{-0.2}$-*BamHI*-GS-mEGFP |
| pET15b mGFP R-MCD$^{0}$ | mGFP-*NdeI*-R-MCD$^{0}$-*BamHI*-TEV-6xHIS |
| pET15b mGFP R-MCD$^{+0.1}$ | mGFP-*NdeI*-R-MCD$^{+0.1}$-*BamHI*-TEV-6xHIS |
| pET15b mGFP R-MCD$^{+0.2}$ | mGFP-*NdeI*-R-MCD$^{+0.2}$-*BamHI*-TEV-6xHIS |
| pmEGFP-N1 R5D5 | CMV(*ΔNdeI*)...*SalI*-*NdeI*-R5D5$^{10}$-*BamHI*-GS-mEGFP |

| | |
|---|---|
| pmEGFP-N1 R7D7 | CMV(*ΔNdeI*)...*SalI*-*NdeI*-R7D7[7]-*BamHI*-GS-mEGFP |
| pmEGFP-N1 R10D10 | CMV(*ΔNdeI*)...*SalI*-*NdeI*-R10D10[5]-*BamHI*-GS-mEGFP |
| pcDNA3.1+ RED0-mCherry | pcDNA3.1+…*NheI*-mCherry-GSGSG-RED$^{0}$-Stop-*XbaI* |
| pcDNA3.1+ RED0.1-mCherry | pcDNA3.1+…*NheI*-mCherry-GSGSG-RED$^{+0.1}$-Stop-*XbaI* |
| pcDNA3.1+ RED0.2-mCherry | pcDNA3.1+…*NheI*-mCherry-GSGSG-RED$^{+0.2}$-Stop-*XbaI* |

**Sequence accession numbers**

| Protein | Uniprot accession | Isoform |
|---|---|---|
| **SPA-5** | Q7RXW5 | NCU01466 |
| **SRSF1** | Q07955 | NP_008855.1 |
| **U1-70k/snRNP70** | P08621 | NP_003080.2 |
| **CPSF6** | Q16630 | NP_001287876.1 |
| **NELF-E** | P18615 | NP_002895.3 |
| **FIP1L1** | Q6UN15 | NP_112179.2 |
| **NCOR1** | O75376 | NP_006302.2 |
| **RERE** | Q9P2R6 | NP_036234.3 |
| **UPF2** | Q9HAU5 | NP_542166.1 |
| **INO80** | Q9ULG1 | NP_060023.1 |
| **SRSF3** | P84103 | NP_003008.1 |
| **SRSF2** | Q01130 | NP_003007.2 |
